# ResQ, a release factor-dependent ribosome rescue factor in the Gram-positive bacterium *Bacillus subtilis*

**DOI:** 10.1101/732420

**Authors:** Naomi Shimokawa-Chiba, Claudia Müller, Keigo Fujiwara, Bertrand Beckert, Koreaki Ito, Daniel N. Wilson, Shinobu Chiba

**Author notes:** These authors contributed equally (Naomi Shimokawa-Chiba, Claudia Müller). Correspondance (S.C.), (D.N.W.).

## Abstract

Rescue of the ribosomes from dead-end translation complexes, such as those on truncated (non-stop) mRNA, is essential for the cell. Whereas bacteria use *trans*-translation for ribosome rescue, some Gram-negative species possess alternative and release factor (RF)-dependent rescue factors, which enable an RF to catalyze stop codon-independent polypeptide release. We now discover that the Gram-positive *Bacillus subtilis* has an evolutionarily distinct ribosome rescue factor named ResQ. Genetic analysis shows that *B. subtilis* requires the function of either *trans*-translation or ResQ for growth, even in the absence of proteotoxic stresses. Biochemical and cryo-EM characterization demonstrates that ResQ binds to non-stop stalled ribosomes, recruits homologous RF2, but not RF1, and induces its transition into an open active conformation. Although ResQ is distinct from *E. coli* ArfA, they use convergent strategies in terms of mode of action and expression regulation, indicating that many bacteria may have evolved as yet unidentified ribosome rescue systems.

## Introduction

Faithful translation requires accurate initiation, elongation and termination. In translation termination, the stop codon situated in the A-site of the ribosome recruits a release factor (RF), which then hydrolyzes the peptidyl-tRNA ester bond to release the polypeptide product from the ribosome. In bacteria, RF1 recognizes UAA and UAG while RF2 recognizes UAA and UGA through their PxT and SPF codon recognition motifs, respectively^1^. These RFs contain the hydrolysis active site motif, GGQ, for catalysis. Polypeptide release is then followed by dissociation of the ribosome from mRNA into the small and large subunits by a processes mediated by ribosome recycling factor (RRF) and elongation factor G (EF-G)^2^.

However, the termination-recycling event can be perturbed when mRNA has aberrant features, one of which is the absence of an in-frame stop codon. The mRNA lacking a stop codon, called a non-stop mRNA, causes stalling of the ribosome at the 3’ end because recruitment of RFs to the ribosome requires the interaction of a stop codon recognition motif of the RF with a cognate stop codon. Since the role of termination is not only to define the end of the protein, but also to recycle the ribosome for the next round of translation initiation, a failure in termination lowers the cellular capacity of protein synthesis, unless dealt with by the cellular quality control mechanisms. Indeed, a loss of function in the quality control machinery leads to an accumulation of dead-end translation products, which was estimated to represent ~2-4% of the translation products in *E. coli*^3^, and results in lethality^4,5^.

Living organisms have evolved mechanisms that resolve non-productive translation complexes produced by ribosome stalling on non-stop mRNAs. Such quality control is also called ribosome rescue. In eukaryotic cells, the Dom34/Hbs1 complex, together with Rli1/ABCE1, mediates ribosome rescue on truncated mRNAs^4,6,7^. In bacteria, two distinct mechanisms operate in the resolution of non-stop nascent chain-ribosome complexes, *trans*-translation and stop codon-independent peptide release from the ribosome^4,5,8,9^. The latter mechanism can further be classified into two classes, RF-dependent and RF-independent. The crucial player in *trans*-translation is the *transfer*-messenger RNA (tmRNA), which is encoded by *ssrA*. tmRNA cooperates with SmpB, which mediates ribosomal accommodation of tmRNA at the ribosomal A-site^10,11^. The tmRNA is composed of tRNA- and mRNA-like domains. The former can be charged with alanine, which then accepts the non-stop peptide and is elongated further according to the mRNA-like coding function of tmRNA until the built-in stop codon is reached. The result is the formation of the non-stop polypeptide bearing an extra *ssrA*-encoded sequence (15 amino acids in *B. subtilis*) and dissociation of the ribosome from the non-stop mRNA. The SsrA tag sequence promotes proteolytic elimination of the non-stop polypeptide via targeting to cellular proteases. *Trans*-translation is essential for the growth of some bacteria^5,12^. The essentiality lies in the liberation of the ribosome from non-stop mRNA, but not in proteolytic degradation of the translation products^13,14^. Bacterial species that can survive without *trans*-translation often possess one or more alternative ribosome rescue factor(s), such as ArfA, ArfB, and ArfT, which are involved in stop codon-independent cleavage of the non-stop peptidyl-tRNA^5^.

ArfA was identified in *E. coli* by a genetic screening for a mutation showing synthetic lethality with the loss of *ssrA*^15^. ArfA is an RF-dependent ribosome rescue factor, which recruits RF2, but not RF1, to the non-stop stalled ribosome complexes to induce hydrolysis of the dead-end peptidyl-tRNA^16^. Interestingly, ArfA itself is produced from a non-stop mRNA, such that it is strongly down-regulated by *trans*-translation in wild-type cells and only induced significantly upon dysfunction of *trans*-translation. Thus, it has been suggested that tmRNA-SmpB is the primary rescue factor, and the ArfA-RF2 system serves as a back-up system^17,18^. ArfB (YaeJ), identified as a multicopy suppressor of the *ssrA/arfA* double mutant^19,20^, contains its own GGQ catalytic motif, enabling it to act as an RF-independent ribosome rescue factor^21^. Although the physiological role of ArfB in *E. coli* is unknown, its homologs are widely distributed among both Gram-positive and -negative bacteria^5,20,22^, as well as eukaryotic mitochondria^23^. ArfT in *Francisella tularensis*, a member of γ-proteobacteria that lacks both ArfA and ArfB homologs^24^, is essential in the absence of tmRNA. Like ArfA, ArfT is an RF-dependent ribosome rescue factor, although ArfT can function with either RF1 or RF2. Phylogenetic distribution of ArfA is limited to a subset of β- and γ-proteobacteria whereas that of ArfT is limited to a subset of γ-proteobacteria. To date, RF-dependent ribosome rescue factors have only been reported in Gram-negative bacteria.

*Bacillus subtilis*, a Gram-positive bacterium, can also survive without *ssrA*^25^, but no alternative factor for ribosome rescue has been reported in this organism, raising the question of whether the ribosome rescue function is non-essential or alternative factors have escaped identification in *B. subtilis*. In this study, we have addressed this question and identified ResQ (<u>Re</u>scue of the <u>S</u>talled ribosome for <u>Q</u>uality control; formerly YqkK) as a ribosome rescue factor. ResQ has no obvious sequence similarity to other Arf proteins. We show that ResQ is an RF2-dependent ribosome rescue factor, which induces hydrolysis of peptidyl-tRNA in non-stop translation complexes. ResQ is produced naturally from a non-stop mRNA and is thus negatively regulated by *trans*-translation, revealing a conceptually similar regulatory crosstalk as documented for *E. coli* ArfA. Lastly, using single-particle cryo-electron microscopy (cryo-EM), we reveal how ResQ recognizes the presence of truncated mRNAs and recruits and stabilizes an open conformation of RF2 to rescue the stalled ribosomes. ResQ uses the mechanism that is similar but distinct from ArfA. Collectively, our findings lead us to suggest that Gram-positive and -negative bacteria have independently acquired their own unique RF-dependent ribosome rescue systems equipped with a convergent scheme of regulation.

## Results

### ResQ (YqkK) is indispensable for the growth of *trans*-translation-deficient *B. subtilis*

The dispensability of *trans*-translation in *B. subtilis* raises the possibility that it contains an alternative ribosome rescue factor. With the reasoning that the loss of such a factor would make the bacterial growth dependent on *trans*-translation proficiency, we searched for the chromosomal gene knockouts that cause synthetic lethality with the deficiency of *trans*-translation. We used a strain with chromosomal deletion of *smpB*, which encodes the *trans*-translation co-factor but having a plasmid carrying the wild-type *smpB* as well as *lacZ* genes (**Fig. 1a**). This rescue plasmid was a derivative of pLOSS*^26^ driven by a temperature sensitive (Ts) replicon, such that it is lost frequently at high temperature. We prepared chromosomal DNA from a mixture of the BKE library strains, a collection of mutants individually disrupted for the 3,968 non-essential *B. subtilis* genes by replacements with the erythromycin resistance marker (*ery*)^27^, and used it to transform (by homologous recombination) the strain for the screening described above. Transformant mixtures were then incubated at 50°C to destabilize the rescue plasmid and plated on selective agar containing X-Gal (see materials and methods). Whereas clones that did not depend on SmpB had segregated out the plasmid and formed white colonies, those requiring SmpB survived only when they had retained the plasmid and formed blue colonies due to the plasmid-encoded β-galactosidase (**Fig. 1a**). Among ~74,000 transformants, we picked up 42 blue colonies and determined the chromosomal locations of the *ery* inserts by DNA sequencing, followed by elimination of false-positive clones by retransformation experiments. These procedures left clones with an *ery* disruption of *yqkK* (renamed *resQ*, see below), which makes SmpB indispensable for survival.

**Fig. 1.**
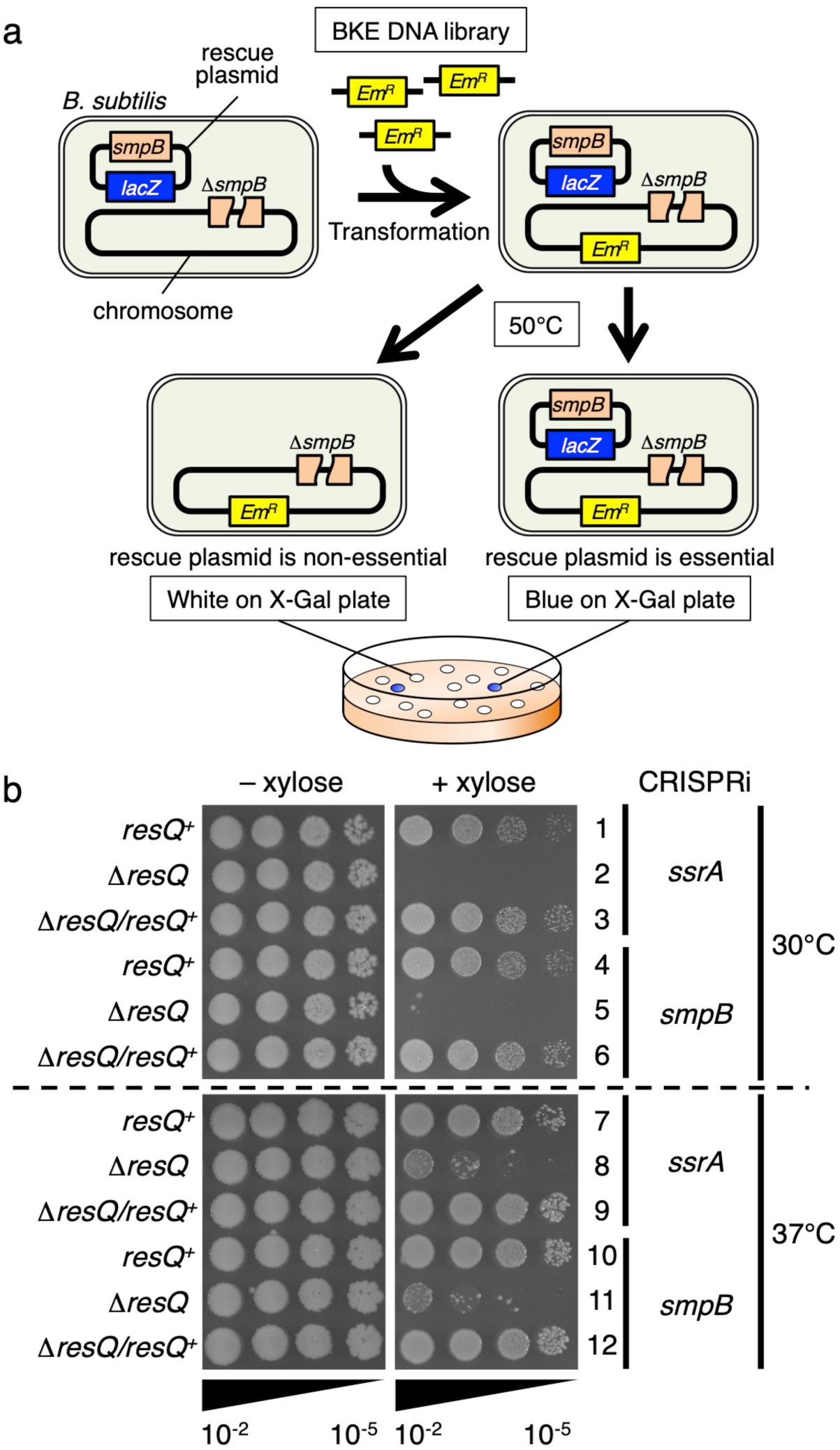
Simultaneous inactivation of ResQ and *trans*-translation leads to synthetic growth defects. **a**, A schematic representation of the synthetic lethal screening to identify genes whose absence causes synthetic lethal phenotype with the deficiency of *trans*-translation. **b**, Xylose-inducible CRISPRi was targeted to *ssrA* (lines 1-3, 7-9) or *smpB* (lines 4-6, 7-9) in the *B. subtilis* strains indicated at the left by the genotypes of the *resQ* gene (Δ*resQ/resQ*^+^ signifies the presence of *resQ*^+^ in an ectopic locus). Cultures prepared in the absence of xylose were serially diluted (from 10^−2^ to 10^−5^) and spotted onto LB agar plates with or without 1% xylose, as indicated at the top, for incubation at 30°C (upper) or 37°C (lower) for 17 hours.

We validated the growth requirement features of *B. subtilis* for ResQ and the *trans*-translation system by an independent approach. We used CRISPR interference (CRISPRi)^28^ to conditionally silence either *ssrA* or *smpB* (**Fig. 1b**). To do this, the catalytically inactive and xylose-inducible variant of Cas9 (dCas9) was integrated into the chromosomal *lacA* locus of the wild type and the Δ*resQ* strains. In addition, the constitutively expressed small guide RNA (sgRNA) that targets either *ssrA* or *smpB* was integrated into the *amyE* locus of the same strains. All the bacteria grew normally when dCas9 was uninduced in the absence of xylose (**Fig. 1b**, left panels). Growth of the *resQ*^+^ cells was not affected by 1% xylose, which induced dCas9 to silence *ssrA* or *smpB* (lanes 1, 4, 7 and 10, right panels). By contrast, growth of the Δ*resQ* mutant was severely impaired in the presence of xylose, which led to CRISPRi-mediated repression of *ssrA* or *smpB* (lanes 2, 5, 8 and 11, right panels). Expression of *resQ^+^* from an ectopic locus restored the growth defect associated with the *trans*-translation deficiency, substantiating that ResQ is responsible factor for the synthetic growth phenotype (lanes 3, 6, 9 and 12). These results indicate that ResQ is required for optimal growth in the absence of sufficient activity of *trans*-translation in *B. subtilis*.

### ResQ is expressed from a non-stop mRNA due to internal transcription termination and regulated negatively by *trans*-translation

The *resQ* gene contains 71 sense codons followed by a stop codon. We note that it contains a typical rho-independent transcription terminator sequence within the coding region, raising an intriguing possibility that ResQ is translated naturally from a non-stop mRNA lacking the 3’ region including the stop codon (**Fig. 2a**). If this were the case, the translation product should initially consist of approximately 62 amino acids in the form of peptidyl-tRNA. However, translation of such mRNAs is likely dealt with by the *trans*-translation mechanism of ribosome rescue, which would add the SsrA-tag sequence of 15 amino acids to the ResQ non-stop product. It follows then that the ResQ product would be rapidly degraded by cellular proteases, such that the protein level should be strongly down-regulated in *trans*-translation proficient cells.

**Fig. 2.**
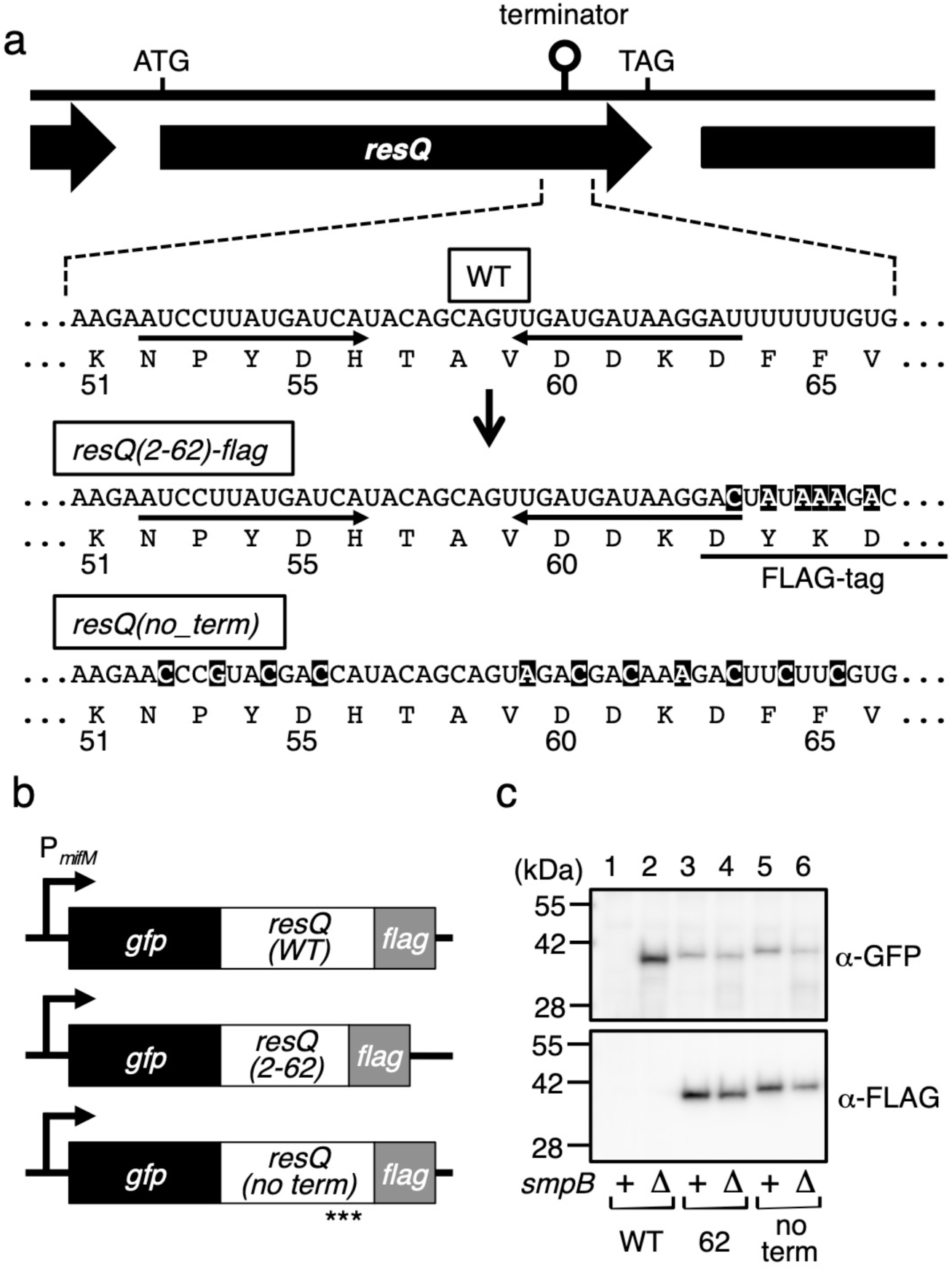
The cellular abundance of ResQ is negatively regulated by *trans*-translation. **a**, A schematic representation of the *resQ* open reading frame, showing the intrinsic transcriptional terminator sequence within the coding region (indicated by “terminator” at the top and the WT enlarged view below). Shown also are the corresponding sequences of the *gfp-resQ(2-62)-flag* and *gfp-resQ(no_term)-flag* constructs (see b), in which nucleotide substitutions and the FLAG tag are indicated by reverse and underline, respectively. **b**, Schematic representations of the *gfp-resQ-flag* constructs with wild type terminator sequence (*gfp-resQ(2-71)-flag*) and its derivatives with defective terminator signals. These *gfp-resQ* derivatives were placed under the constitutive *mifM* promoter. Asterisks indicate synonymous mutations. **c**, Cellular accumulation of the products of the wild type construct (lanes 1, 2) as well as the *resQ(2-62)* (lanes 3, 4) and the *resQ(no_term)* (lanes 5, 6) constructs. They were expressed in the *smpB*^+^ (odd numbers) or the Δ*smpB* (even numbers) strains and analyzed by anti-GFP (upper) or anti-FLAG (lower) immunoblotting.

To test this possibility, we examined the impact of *trans*-translation on the accumulation of a series of ResQ-derived proteins. We constructed a gene fusion consisting of the coding sequences for green fluorescence protein (GFP), ResQ (full length) and FLAG that are connected in-frame in this order from the N-terminus to the C-terminus (**Fig. 2b**). Anti-GFP immunoblotting was used to detect products related to GFP-ResQ-FLAG, when it was expressed in the wild type and the Δ*smpB* cells. Strikingly, the wild-type cells did not produce any protein detectable with the GFP antibodies (**Fig. 2c**, lane 1, upper panel). By contrast, the Δ*smpB* cells produced a product that formed an intense band slightly below the 42 kDa marker (**Fig. 2c**, lane 2, upper panel). This species of protein did not react with anti-FLAG (lane 2, lower panel), indicating that it lacked the C-terminal region. These results are consistent with the notion that transcription termination within the ResQ-coding region generates a truncated mRNA that ends within *resQ*. The translation products are SsrA-tagged and degraded rapidly in the wild type strain, while they accumulate in the *trans*-translation-deficient Δ*smpB* cells.

To examine the above scenario further, we disrupted the transcription terminator either by internal deletion or by synonymous substitutions (**Fig. 2a,b**). In the former, we deleted the codons 63-71 of *resQ* in the context of GFP-ResQ-FLAG to eliminate the 3’ T stretch of the transcriptional terminator (GFP-ResQ62-FLAG). In the latter, we introduced synonymous mutations to disrupt the secondary structure required for termination (GFP-ResQ(no_term)-FLAG) (**Fig. 2a,b**). Both of these terminator-less mutant forms of GFP-ResQ-FLAG accumulated equally in the wildtype and Δ*smpB* strains as a form reactive with both anti-GFP and anti-FLAG (**Fig. 2c**, lanes 3-6). Thus, the terminator sequence indeed functions to prevent the expression of the downstream coding sequence. Also, the terminator is a major element that triggers the ResQ down-regulation by the SsrA tag-dependent degradation.

These results show that *B. subtilis* is equipped with at least two layers of ribosome rescue mechanisms, *trans*-translation and ResQ-dependent peptidyl-tRNA hydrolysis (see below). The *trans*-translation-dependent down-regulation indicates that ResQ is the secondary ribosome-rescue factor that is only produced upon dysfunction of *trans*-translation, the primary ribosome rescue mechanism. Thus, the internal transcription terminator in ResQ provides the means for this bacterium to accomplish the compensatory and vectorial regulation for the maintenance of ribosome rescue capability. In this context, ResQ bears a striking similarity to the *E. coli* alternative rescue factor, ArfA, which is also synthesized from a non-stop mRNA^17,18^.

### ResQ recruits RF2 to hydrolyze non-stop peptidyl-tRNAs

We characterized ResQ biochemically by examining whether it could induce polypeptide release, as expected for an alternative ribosome rescue factor. To do this, we purified ResQ in the form of ResQ62-His_6_, which lacks the C-terminal 9 amino acid residues encoded by the *resQ* gene but not by the *resQ* mRNA (see above); a hexahistidine (His_6_) tag was attached to the C-terminus to aid purification. We considered that the absence of the C-terminal 9 amino acids was physiological. Also, the absence of the 3’ coding region of *resQ* should disrupt the internal transcription terminator signal and enable the attachment of the His_6_ tag. Since ResQ lacks a GGQ motif critical for peptidyl-tRNA hydrolysis, we hypothesized that ResQ requires a release factor (RF) to hydrolyze peptidyl-tRNA. Therefore, we also purified *B. subtilis* RF1 and RF2. For in vitro translation with defined translation components, we used the *Bs* hybrid PURE system^29^, a modified version of the PURE coupled transcription-translation system^30^, in which the original *E. coli* ribosomes were replaced with *B. subtilis* ribosomes. We omitted RFs in the *Bs* hybrid PURE system unless otherwise stated.

We used DNA fragments encoding GFP but without an in-frame stop codon (GFP-ns) to direct in vitro transcription and translation with *Bs* hybrid PURE system and separated the translation products by neutral pH SDS-PAGE, which preserved the peptidyl-tRNA ester bond. The major product, migrating between the 42 kDa and the 55 kDa markers, represents the peptidyl-tRNA (GFP-tRNA; **Fig. 3a**, lane 1), as treatment of the sample with RNase A before electrophoresis down-shifted this band to the position near the 28 kDa marker, indicative of tRNA removal (GFP; lane 2). Thus, the non-stop template indeed produced a translation-arrested state of ribosome-nascent chain complex, which was unaffected when the reaction mixture included *B. subtilis* RF1 or RF2 (lanes 3-6), as expected from the absence of a stop codon in the template. We then addressed the effects of ResQ. Neither ResQ by itself, nor its combination with *B. subtilis* RF1, affected the production of GFP-tRNA (lanes 7-10). By contrast, the addition of both ResQ and *B. subtilis* RF2 to the *Bs* hybrid PURE system resulted in the production of the hydrolyzed GFP band with a concomitant decrease in the level of GFP-tRNA (lanes 11 and 12). Thus, ResQ and RF2 cooperatively catalyze peptide release from the non-stop stalled ribosome.

**Fig. 3.**
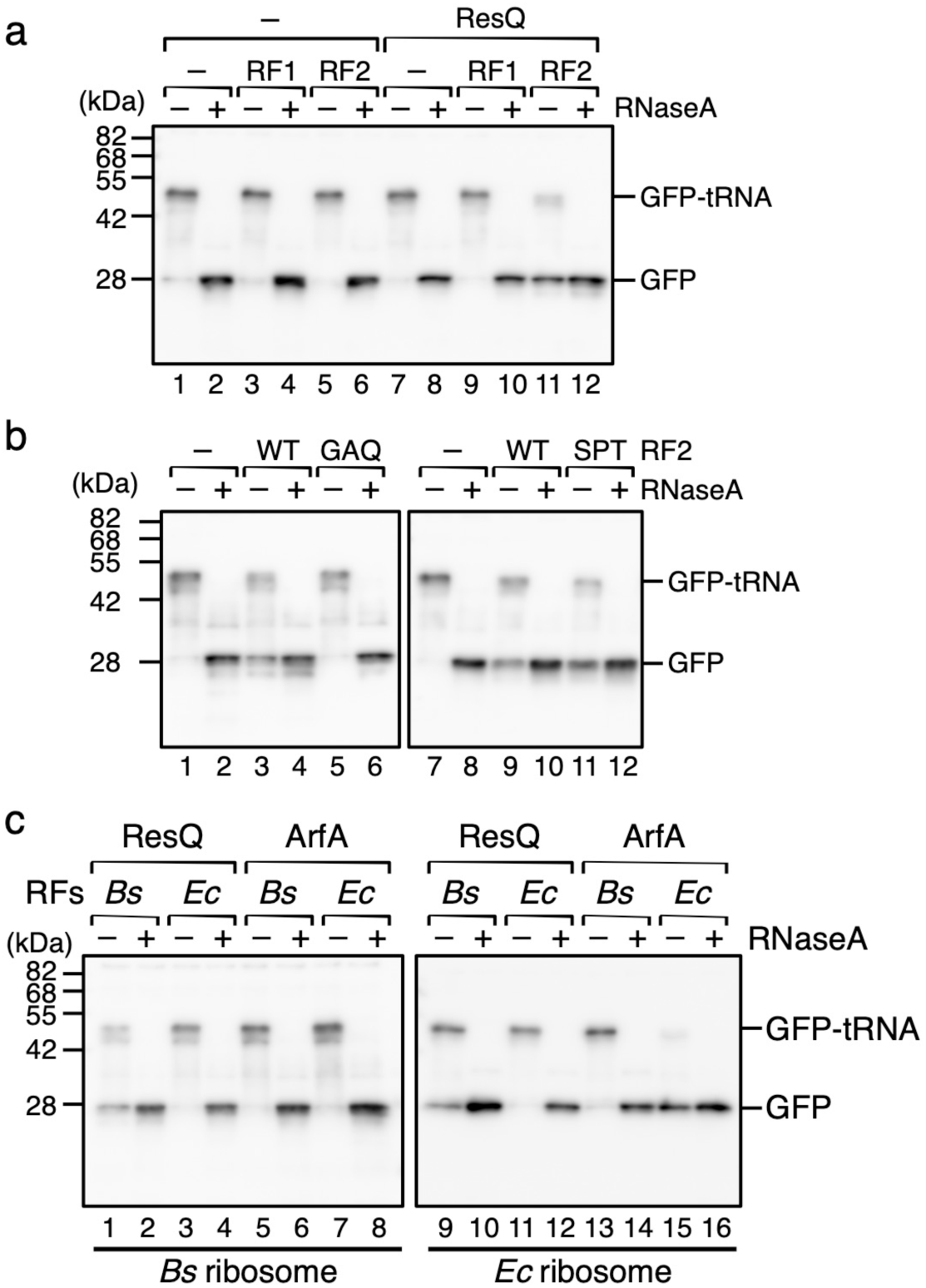
*In vitro* activity of ResQ to hydrolyze non-stop peptidyl-tRNA. **a**, ResQ in combination with RF2 cleaves the GFP-tRNA non-stop translation product. *In vitro* translation using *Bs* hybrid PURE system was directed by the *gfp-ns* template. The reaction mixtures contained purified ResQ62-His_6_ (lanes 7-12), purified *B. subtilis* RF1 (lanes 3, 4, 9, 10) and RF2 (lanes 5, 6, 11, 12), as indicated. Translation was allowed to proceed at 37°C for 20 min, and the products were divided into two parts, one of which was treated with RNaseA, as indicated. Samples were then analyzed by SDS-PAGE under neutral pH conditions, followed by anti-GFP immunoblotting. **b**, ResQ-dependent peptidyl-tRNA hydrolysis activity of RF2 requires its GGQ active site but not SPF stop codon recognition motif. *In vitro* translation using *Bs* hybrid PURE system was directed by the *gfp-ns* template in the presence of combinations of ResQ, wild type RF2 (lanes 3, 4, 9, 10), RF2(GAQ) (lanes 5, 6) and RF(FPT) (lanes 11, 12), as indicated. The translation products were analyzed by anti-GFP immunoblotting as described above. **c**, Interspecies compatibility of the RF-dependent rescue factor functions. *In vitro* translation of *gfp-ns* was carried out using *Bs* hybrid PURE system (lanes 1-8) or *Ec* PURE system (lanes 9-16) in the presence of combinations of purified ResQ (lanes 1-4, 9-12), *E. coli* ArfA (lanes 5-8, 13-16), RFs (RF1 plus RF2) purified from *B. subtilis* (lanes 1, 2, 5, 6, 9, 10, 13, 14) and RFs purified from *E. coli* (lanes 3, 4, 7, 8, 11, 12, 15, 16). The translation products were analyzed by anti-GFP immunoblotting as described above.

### Distinct requirements for ribosome rescue and the canonical termination functions of RF2

Given that ResQ and RF2 cooperatively induce hydrolysis of non-stop peptidyl-tRNA, the role of RF2 would be to execute the catalysis. Consistent with this expectation, a catalytically inactive RF2 variant, RF2(GAQ), whose GGQ active site had been mutated to GAQ, no longer stimulated hydrolysis of GFP-tRNA even in the presence of ResQ (**Fig. 3b**, lanes 5 and 6). RF2 possesses the conserved SPF motif as an essential element for the stop codon recognition in termination^31^. We addressed whether this motif is required for the ribosome rescue function of RF2 by mutating it to SPT, which abolishes the termination activity^31^. In the *in vitro* peptidyl-tRNA hydrolysis assay, the RF2(SPT) was as active as the wild-type RF2 in the ResQ-dependent cleavage of GFP-tRNA (**Fig. 3b**, lanes 9-12), in comparison with the parallel control reaction without RF2 (lanes 7 and 8). These results demonstrate that the stop codon recognition motif of RF2 is dispensable for the ribosome rescue function.

Ribosome stalling can also be induced by specific amino acid sequences of the nascent polypeptides for regulatory purposes. Such regulatory nascent polypeptides include *E. coli* SecM, *B. subtilis* MifM and *Vibrio alginolyticus* VemP^32,33^. Ribosome stalling in these cases needs to be adequately regulated, such that it is subject to conditional and specific mechanisms of cancellation^34,35^. Unregulated rescue could be counterproductive in these cases. Here we could show that the *B. subtilis* MifM stalling is refractory to both the *trans*-translation as well as ResQ ribosome rescue mechanisms *in vivo* and *in vitro* (**Supplementary Fig. 1**). Thus, we conclude that ResQ is an RF-dependent ribosome rescue factor in *B. subtilis* that rescues stalling on non-stop mRNAs, but not regulatory stalling peptides, leading us to suggest the renaming of YqkK to ResQ (<u>Re</u>scue of the <u>S</u>talled ribosome for <u>Q</u>uality control of translation). ResQ is the first reported example of an RF-dependent ribosome rescue factor in a Gram-positive bacterium.

### Low interspecies compatibility of the RF-dependent ribosome rescue systems

ResQ homologs were conserved among a subset of *Bacillaceae* family members, mainly among those belonging to the *Bacillus* genus. The distinct phylogenetic distribution and the unique sequence features, which are unrelated to the proteobacterial ArfA or ArfT (**Supplementary Fig. 2**), suggest that ResQ has evolved independently of other alternative rescue factors. Moreover, the narrow phylogenetic distributions of the RF-dependent rescue factors also implies that they emerged relatively late in evolution. Translation components are largely conserved across the species, but they have undergone some micro diversifications. If the alternative rescue factors evolved more recently than the translation factors, their interactions with the translation components, such as the ribosome and an RF, might be species-specific. Indeed, *F. tularensis* ArfT can work with *F. tularensis* RF1 or RF2, but not *E. coli* RFs^24^. Also, *E. coli* ArfA fails to recruit *Thermus thermophilus* RF2^36,37^. We tested the compatibility of *B. subtilis* ResQ and *E. coli* ArfA with heterologous RFs in vitro. While ResQ efficiently hydrolyzed GFP-tRNA in the *Bs* PURE system supplemented with RF1 and RF2 of *B. subtilis* (**Fig. 3c**, lanes 1 and 2), it did not work with the *E. coli* RFs (lanes 3 and 4). Interestingly, ResQ was functional when the substrate was translated by the *E. coli* ribosome, provided that the *B. subtilis* RFs were available (lanes 9-10). These results suggest that ResQ interaction with RF2 is species-specific, but its interaction with the ribosome is rather promiscuous.

By contrast, we could show that *E. coli* ArfA requires both the RF2 as well as the ribosomes to be derived from *E. coli*. Specifically, ArfA did not work if combined with the homologous RF2 when the substrate was translated using *Bs* PURE system (**Fig. 3c**, lanes 7, 8), nor did it work with *B. subtilis* RF2 (lanes 5, 6) if the substrate was translated by *E. coli* ribosomes (lanes 13 and 14). The *E. coli* ArfA-mediated hydrolysis of GFP-tRNA was only observed when the ribosomes and RFs were both derived from *E. coli* (lanes 15 and 16). Thus, *E. coli* ArfA is incompatible with the *B. subtilis* ribosomes and RF2, analogous to that observed previously between *E. coli* ArfA and *T. thermophilus* RF2^36,37^. The high specificity of molecular interactions involving the RF-dependent rescue factors is in contrast to the broader interactions in the tmRNA- and ArfB-based rescue pathways (see Discussion).

### Cryo-EM structure of a ResQ-RF2-non-stop-ribosome complex

Since *B. subtilis* ResQ and RF2 can rescue *E. coli* ribosomes stalled on truncated non-stop mRNAs (**Fig. 3c** and **Supplementary Fig. 3**), we formed ResQ-RF2-non-stop 70S ribosome (ns70S) complexes by incubating *B. subtilis* ResQ and RF2 with *E. coli* ns70S complexes used previously for ArfA^38^. By substituting the wildtype *B. subtilis* RF2 with a catalytically inactive GGP mutant^39,40^, peptidyl-tRNA hydrolysis and therefore recycling of the ns70S complex was prevented (**Supplementary Fig. 3**). Cryo-EM analysis of the ResQ62His-RF2-GGP-ns70S complex (herein referred as ResQ-RF2-ns70S) and extensive *in silico* sorting of this dataset yielded a major subpopulation of ribosomal particles (>80%) that contained stoichiometric occupancy of ResQ, RF2 and P-site tRNA (**Supplementary Fig. 4**). Refinement of this subpopulation led to a final cryo-EM reconstruction of the ResQ-RF2-ns70S (**Fig. 4a**), with an average resolution of 3.15 Å (**Supplementary Fig. 4** and **Supplementary Table 1**). The cryo-EM density for ResQ was well-resolved with local resolution ranging between 3.0-3.6 Å (**Fig. 4b**), enabling residues 2-55 of ResQ to be modelled *de novo* (**Fig. 4c,d**). ResQ contains an N-terminal α-helix α1 (residues 4-17) followed by a short α-helical turn (α2, residues 21-25), as well as a β-strand (β1, residues 35-38) and a short C-terminal α-helix (α3, 40-47) followed by a positively charged region (residues 48-55) (**Fig. 4d**).

**Fig. 4.**
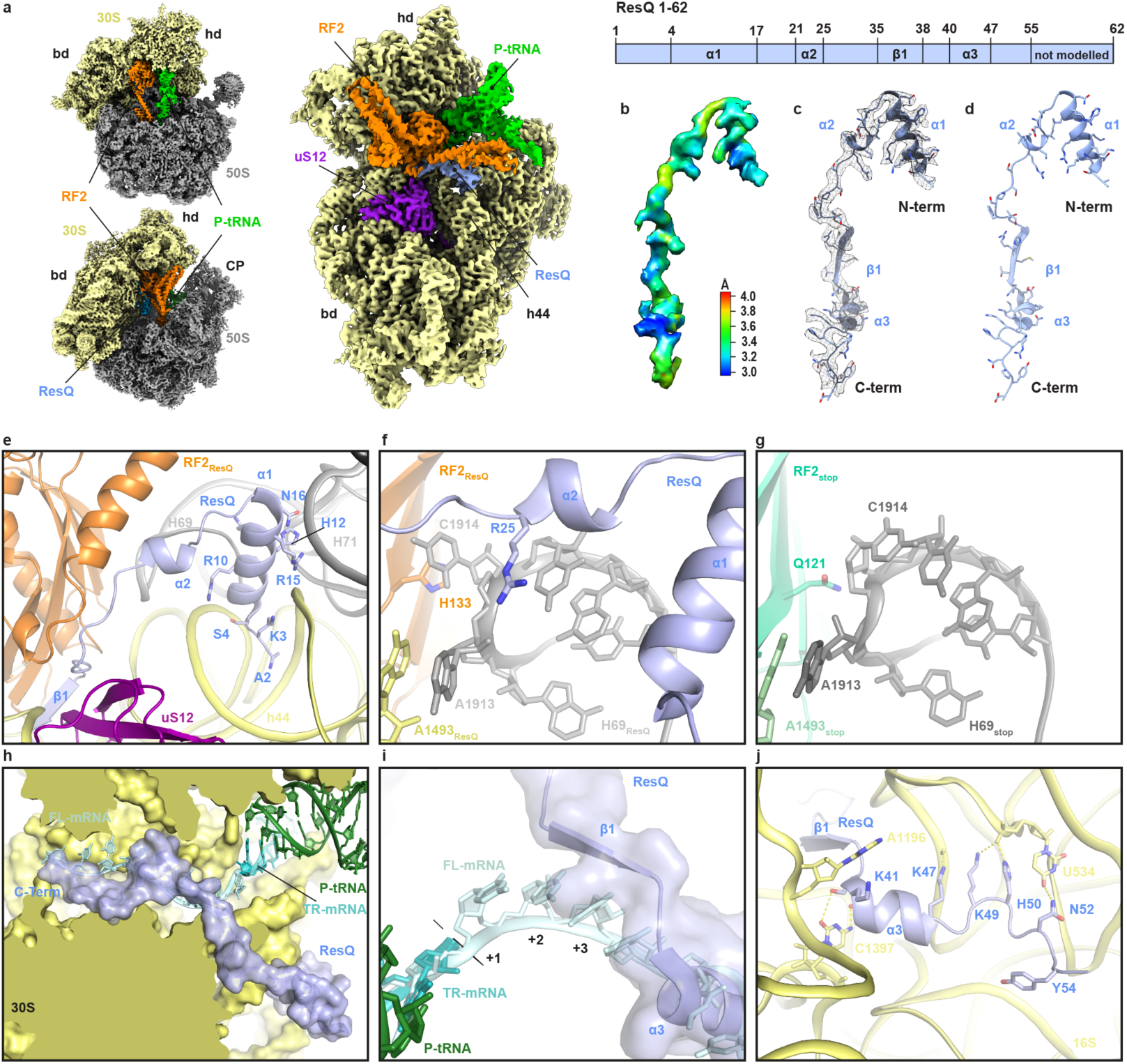
Cryo-EM structure of ResQ-RF2-ns70S complex. **a**, Different views of the cryo-EM map of the ResQ-RF2-ns70S complex with isolated densities highlighting the 30S (yellow; hd, head and bd, body) and 50S (grey; CP, central protuberance) subunits, P-site tRNA (green), RF2 (orange) and ResQ (blue). **b-c**, Isolated electron density for ResQ (b) colored according to local resolution and (c) shown as mesh (grey) with fitted molecular model for ResQ. **d**, Model for ResQ with features highlighted corresponding to the schematic of ResQ protein, including α-helical and β-strand regions. **e**, The N-terminus of ResQ (blue) interacts both h44 of the 16S rRNA (yellow) and H69 and H71 of the 23S rRNA (grey). **f**, The conserved R25 of ResQ (blue) stacks upon U1915 and causes C1914 to flip out and stack upon H133 of RF2. **g**, Same view as (f) but showing the RF2_stop_ (lime) and the conformation of H69 for a canonical termination complex (PDB ID 4V5E^47^). **h**, Transverse section of the 30S subunit (yellow) to reveal the mRNA channel showing a superimposition of full-length mRNA (FL-mRNA, cyan) with truncated non-stop mRNA (TR-mRNA, teal), P-site tRNA (green) and surface representations of ResQ (blue). **i**, Superimposition of FL-mRNA (cyan) with TR-mRNA (teal), P-site tRNA (green) and transparent surface representation of ResQ (blue). The first (+1), second (+2) and third (+3) nucleotides of the A-site codon of the FL-mRNA are indicated. **j**, Interaction of the C-terminus of ResQ (blue) with the 16S rRNA showing potential hydrogen bonds with yellow dashed lines.

### Interaction of ResQ with the non-stop 70S ribosome

The binding site of ResQ is located predominantly on the 30S subunit in the vicinity of the decoding center, where it spans from the top of helix 44 (h44) of the 16S rRNA past the ribosomal protein uS12 and reaches into the mRNA channel formed by the head and body of the 30S (**Fig. 4a**). The overall binding site of ResQ on the ribosome (**Fig. 4A**) is similar, but distinct, to that observed previously for *E. coli* ArfA^36–38,41–43^. The N-terminal helix α1 of ResQ resides within the intersubunit space, where highly conserved charged residues (**Supplementary Fig. 2**) establish interactions with the major groove of h44 and the minor groove of H71 of the 23S rRNA (**Fig. 4e**). In addition, Arg25 within helix α2 of ResQ, which is conserved in all ResQ sequences (**Supplementary Fig. 2**), stacks upon U1915 and flips C1914 out of H69, where it stacks upon His133 of RF2 (**Fig. 4f**). This contrasts with the canonical conformation of C1914 within H69 that is observed during translation termination (**Fig. 4g**) as well as ArfA-mediated ribosome rescue. Indeed, these N-terminal α-helices have no counterpart in ArfA, instead the N-terminus of ArfA is unstructured and folds back to interact with uS12^36–38,41–43^ (**Supplementary Fig. 5a-c**).

The C-terminal region of ResQ extends from the decoding center into the mRNA channel and would be incompatible with the presence of a full-length mRNA (**Fig. 4h**), but compatible with a truncated non-stop mRNA (**Fig. 4i**). ResQ exhibits a modest overlap with the second (+2) and third (+3) nucleotide of the A-site codon, but extensive steric clashes would be expected for the subsequent positions (+4 onwards) (**Fig. 4i**), similar to that observed previously for ArfA^36–38,41–43^ (**Supplementary Fig. 5d-i**). Thus, ResQ may also recycle ribosomes stalled on non-stop mRNAs with 1-3 nucleotides extending into the A-site, as shown experimentally for ArfA^44–46^. The positively charged C-terminus of ResQ can form multiple hydrogen bond interactions with 16S rRNA nucleotides that comprise the mRNA channel (**Fig. 4j**). While the interaction network is generally distinct from that observed for ArfA, we note that the mode of contact between the side chains of Lys49 and His50 of ResQ with U534 of the 16S rRNA appears to be shared by ArfA^36–38,41–43^ (**Supplementary Fig. 5j-l**).

### ResQ recruits and stabilizes the open conformation of RF2 on the ribosome

The structure of the ResQ-RF2-ns70S reveals that ResQ recruits RF2 by establishing an extensive interaction surface, specifically encompassing the central portion (residues 30-40) of ResQ and domain 2 (d2) of RF2 (**Fig. 5a**). Similar to ArfA^36–38,41–43^ (**Supplementary Fig. 6A-C**), ResQ also donates the small β-strand (β1) to augment the β-sheet of the superdomain d2/d4 of RF2 (**Fig. 5a**). The overall position of RF2 in ResQ-RF2-ns70S is similar, but slightly shifted, compared to that observed during canonical translation termination^39,47^ (**Fig. 5b** and **Supplementary Fig. 6d-f**)). The shift is larger and more global than reported previously for ArfA^36–38,41–43^ (**Supplementary Fig. 6a-c**), which may arise in part due to the differences between *B. subtilis* and *E. coli* RF2s. The shift affects the loop between the β4-β5 strands of d2 bearing the SPF (*B. subtilis* 202-Ser-Pro-Phe-204) motif, which is involved in the specificity of recognition of the first and second positions of UGA/UAA stop codons^31,39,47^ (**Fig. 5c**). Importantly, the structure illustrates that ResQ, like ArfA, does not interact with the SPF motif and therefore does not directly mimic the presence of a stop codon (**Fig. 5c** and **Supplementary Fig. 6g-i**), which is consistent with our observation that mutations in the SPF motif that impair RF2 termination activity, do not affect ResQ–RF2-mediated ribosome recycling (**Fig. 3b**).

**Fig. 5.**
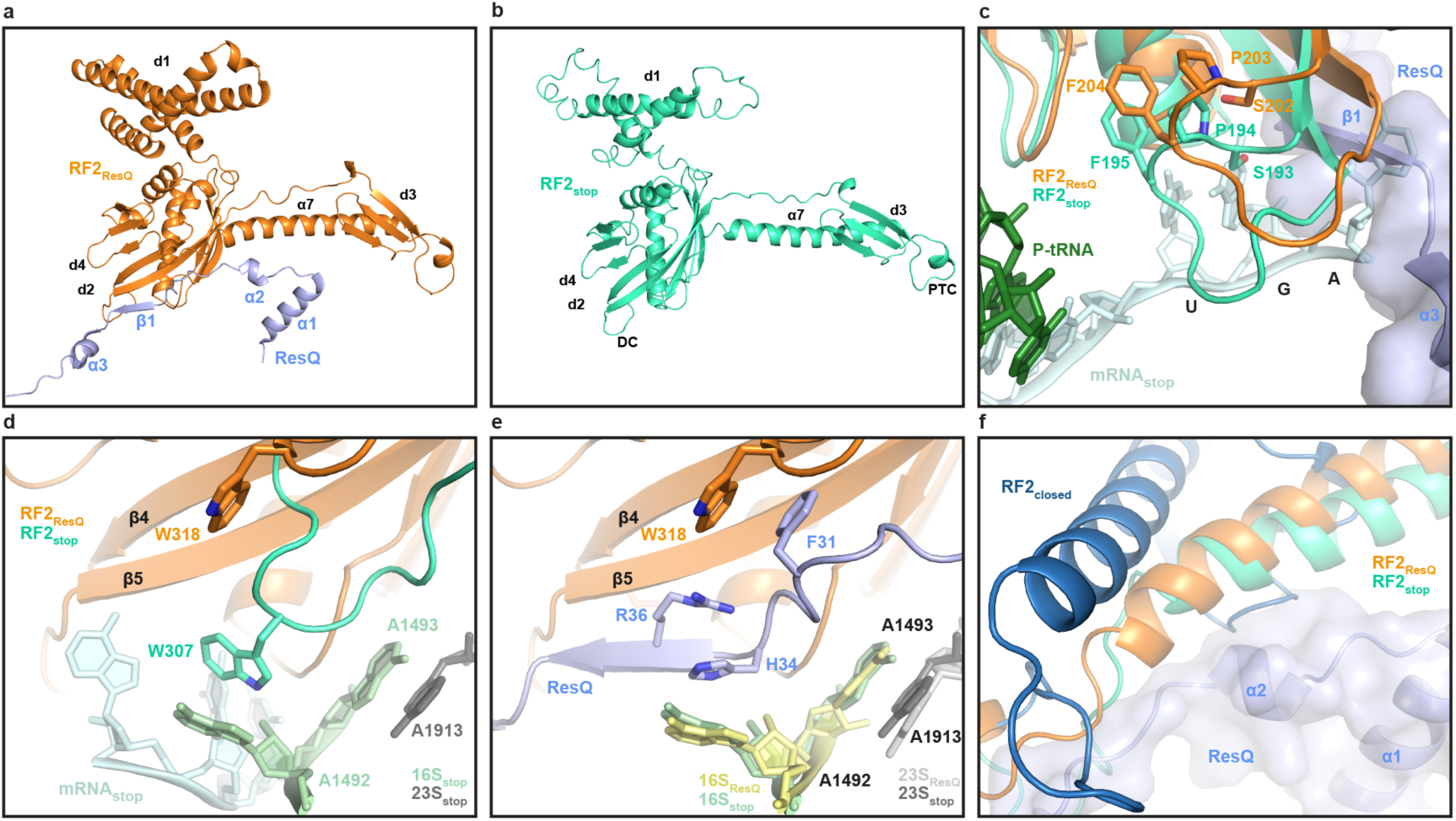
ResQ recruits and stabilizes the open conformation of RF2 on the ribosome. **a-b**, Overview of the interaction of ResQ (blue) with (a) RF2 (RF2_ResQ_, orange) from the ResQ-RF2-ns70S, and with (b) RF2 (RF2_stop_, lime, PDB ID 4V5E) from a canonical termination complex ^47^. RF2 domains 1-4 (d1-d4) and relative positions of the decoding center (DC) and peptidyltransferase center (PTC) are indicated. **c**, Comparison of the relative positions of the SPF motif of RF2_stop_ (lime) and RF2_ResQ_ (orange) with ResQ (blue), P-site tRNA (green) and truncated non-stop mRNA (TR-mRNA, cyan) shown for reference. **d**, Interaction between Trp307 (W307, equivalent to *B. subtilis* Trp318 (W318)) of the switch region of *Thermus thermophilus* RF2_stop_ (lime) and A1492 of the 16S rRNA (green) during decoding of the UGA stop codon of the mRNA (cyan). W318 in the switch loop of *B. subtilis* RF2 (RF2_ResQ_, orange) observed upon ResQ binding is superimposed and arrowed. **e**, Same view as in (d), but showing the conformation of the switch loop of RF2_ResQ_ (orange) and A1492/A1493 (yellow) when ResQ (blue) is present. **f**, Superimposition of the conformation of α-helix α7 of RF2 from the crystal structure of the closed form of RF2 (RF2_closed_; dark blue, PDB ID 1GQE) with RF2_stop_ (lime) and RF2_ResQ_ (orange), with ResQ (blue) shown for reference.

During canonical termination, recognition of the stop codon by RF2 (and RF1) is proposed to stabilize a distinct conformation of the switch loop that directs domain 3 into the PTC ^1,48^. The switch loop conformation is stabilized by stacking interactions between Trp318 (*B. subtilis* numbering) of RF2 and A1492 (h44) as well as between A1493 (h44) and A1913 in H69^39,47^ (**Fig. 5d**). In the ResQ-RF2-ns70S, the presence of ResQ precludes a direct interaction between W318 and A1492 (**Fig. 5e**). Rather, ResQ appears to stabilize a similar conformation of A1492 through stacking interactions with His34 and Arg36 (**Fig. 5e**), whereas a completely distinct conformation of A1492 (and A1493) is adopted in the presence of ArfA (**Supplementary Fig. 7a-c**). Additional stacking interactions are also observed between Phe31 of ResQ and Trp318 within the switch region of RF2 (**Fig. 5e**), which we suggest facilitates the transition from the closed to the open form of RF2 (**Fig. 5f** and **Supplementary Fig. 7d-i**) and thereby enables placement of the GGQ motif within domain 3 (d3) of RF2 at the PTC of the ribosome.

## Discussion

We have shown that Gram-positive bacteria, such as *Bacillus*, possess an RF-dependent ribosome rescue pathway, which had previously been known to occur only in Gram-negative bacteria (**Fig. 6a**). In this pathway in *B. subtilis*, ResQ plays a critical role in the hydrolytic release of the incomplete polypeptide from the non-stop stalled ribosomes. It does so by recruiting RF2 in a stop-codon-independent manner to the otherwise dead-end translation complex, as shown by our biochemical experiments using purified components. Our cryo-EM structure also reveals that ResQ recognizes the empty mRNA channel of a non-stop ribosome complex to recruit and stabilize the active (open) conformation of RF2 on the ribosome (**Fig. 6b**), in a similar but distinct manner to ArfA^36–38,41–43^ (**Fig. 6c**).

**Fig 6.**
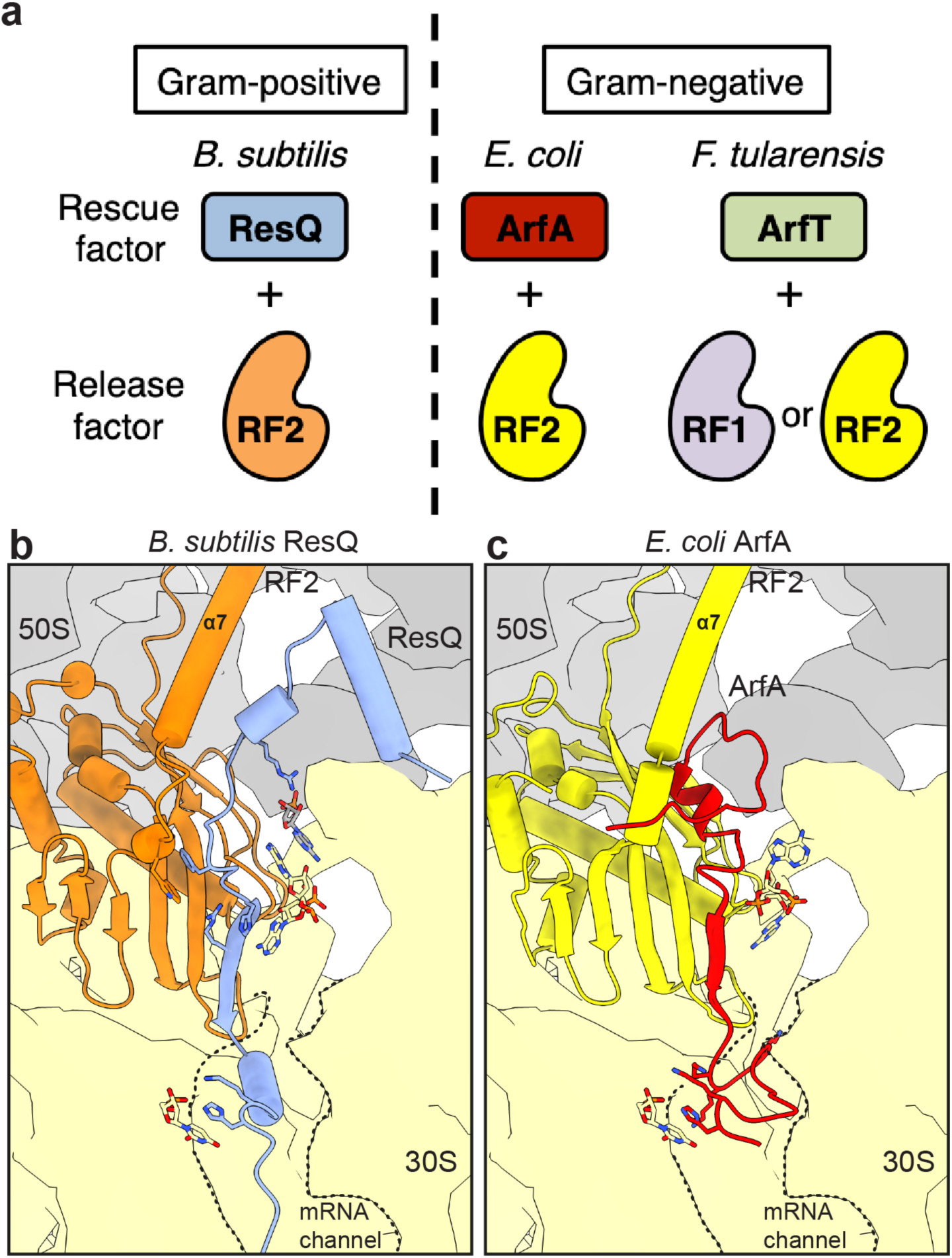
RF-dependent ribosome rescue factors in bacteria. **a**, Independent evolution of the RF-dependent ribosome rescue factors. ArfA and ResQ are unrelated in their amino acid sequences despite the same partner (RF2) specificity. ArfT forms a still distinct group and cooperates with either of RF1 or RF2. They show narrow distributions among bacterial species. **b-c**, A schematic model highlighting the similarities and differences between (b) ResQ- and (c) ArfA-mediated ribosome rescue mechanism.

*In vivo*, the *resQ* deletion mutation exhibits a synthetic lethal phenotype when combined with CRISPRi-mediated knock-down or deletion of either SsrA or SmpB. Thus, ResQ is essential for growth in the absence of *trans*-translation activity. These findings answer the long-standing question of why the *trans*-translation system is not essential in *B. subtilis*, which lacks ArfA, ArfB and ArfT. Instead, ResQ is the alternative ribosome rescue factor in this organism, as well as some other *Bacillus* species. The growth requirement of *B. subtilis* for ResQ and *trans*-translation is observed under “normal” growth conditions without the imposition of proteotoxic stresses. Thus, clearing of constitutively produced aberrant states of translation might be critical for cell survival, and ResQ and *trans*-translation components are the major players in this basic quality control. In this regard, the recently reported RqcH pathway, which adds a protein-degrading polyalanine sequence to the arrested polypeptide on the split 50S subunit of the ribosome^49^, appears to play a minor role in ribosome rescue under normal growth conditions, since RqcH is dispensable for *trans*-translation deficient strains to grow in the absence of stress. By contrast, RqcH appears to be required under more harsh conditions, such as high temperature and the presence of translation-disturbing drugs^49^.

During ResQ-mediated ribosome rescue, peptidyl-tRNA hydrolysis depends on an intact GGQ motif of RF2, indicating that RF2 plays the catalytic role. The role of ResQ is rather to recruit RF2 to the ns70S complex to initiate the events of stop codon-independent translation termination and thereby allow the subsequent ribosome recycling process. We used the *Bs* PURE system containing only a minimal set of essential translation components from *E. coli* together with *B. subtilis* ribosomes to recapitulate the rescue reaction *in vitro*, making it unlikely that other unknown cellular factors, especially those from *B. subtilis*, are required for the process. Moreover, we could show structurally that ResQ alone is sufficient to recruit and induce an active (open) conformation of *Bs*RF2 on the non-stop ribosome complex. Taken together, we propose that ResQ is an RF-dependent ribosome rescue factor in Gram-positive bacteria, such as *Bacillus* (**Fig. 6a**).

In the process of canonical translation termination, stop codon recognition by the SPF motif of RF2 is a prerequisite for the process in which RF2 is accommodated into the A-site of the ribosome and adopting a catalytically active (open) conformation where the GGQ motif is directed into the PTC^39,47,50,51^. However, our structural data show that ResQ does not directly mimic the stop codon in the A-site (**Fig. 5C**), consistent with our observations that the stop codon recognition motif SPF is not required for ribosome rescue (**Fig. 3b**; see ref.^16^ for the similar situation in *E. coli* ArfA). Instead, the role of ResQ is to induce an open conformation of RF2, despite the absence of a stop codon.

ResQ appears to facilitate ribosome rescue using a different mechanism than reported previously for ArfA^36–38,41–43^. For example, during stop codon decoding, the switch loop Trp stacks directly upon A1492 of the 16S rRNA (**Fig. 5d** and **Supplementary Fig. 7a**), whereas during ResQ-mediated recycling, a series of stacking interactions appear to indirectly relay this information - specifically, Phe31 of ResQ stacks upon the switch loop Trp318 of RF2, while His34 (and Arg36) of ResQ stacks upon A1492 (**Fig. 5e** and **Supplementary 7b**). By contrast, during ArfA-mediated rescue, the equivalent Trp residue inserts into a hydrophobic pocket created by ArfA and A1492 adopts a completely unrelated conformation (**Supplementary Fig. 7c**)^36–38,41–43^.

Perhaps the most dramatic difference between ResQ and ArfA is found in the N-terminal region, where ResQ contains two short α-helices that establish multiple interactions with H69 and H71 of the 23S rRNA, whereas the N-terminus of ArfA is unstructured and does not establish any interactions with the 50S subunit (**Supplementary Fig. 5a-c**). Despite this lack of sequence and structural homology, there are some common features between the two rescue systems: Firstly, both ArfA and ResQ (as well as SmpB and ArfB) appear to utilize positively charged C-terminal extensions to interact with the negatively charged 16S rRNA comprising the mRNA channel. Although the details of the interactions are unrelated, we identified a Lys-His (K49-H50) motif that is conserved between ResQ and ArfA, which is used to establish contacts to the backbone of nucleotide U534 of the 16S rRNA (**Supplementary Fig. 5j-l**). Secondly, both ArfA and ResQ contain short ß-strands that augment the ß-sheet in domain 2/4 of RF2, which we presume is important to recruit RF2 to the ribosome (**Supplementary Fig. 5a-c**).

Here we show that ResQ works with RF2, but not RF1, reminiscent of the partner selectivity described previously for ArfA^16^. Analysis of the contacts between ResQ and RF2 within the ResQ-RF2-ns70S complex, and comparison with the sequence alignments between *B. subtilis* RF2 and RF1, indicated that there are two main regions in the RFs that are likely to be responsible for the selectivity of ResQ (**Supplementary Fig. 8**). These encompass residues within the ß-sheet of domain 2/4 of RF2 that are in proximity of Phe31 of ResQ. For example, in RF2 residues 197 and 216 are generally Val and Phe, respectively, whereas in RF1 the equivalent residues are replaced with the smaller Ala and Thr residues, respectively (**Supplementary Fig. 8a**). More dramatic, however, is the lack of sequence conservation between the switch loops of *B. subtilis* RF1 and RF2. In fact, the switch loop of RF1 is one residue longer than in RF2 (**Supplementary Fig. 8b,d**). We also note that the RF-dependent rescue factors show low interspecies compatibility (**Fig. 3c**). The incompatibility of ResQ to work with *E. coli* RF2 is not surprising given the relatively low sequence conservation observed within the switch region (**Supplementary Fig. 8d**). Moreover, sequence differences are also observed with α-helix 7 between *B. subtilis* and *E. coli* RF2 that could contribute to the interactions with ResQ (**Supplementary Fig. 8c).** Collectively, these differences suggest that even if ResQ could recruit *B. subtilis* RF1 or *E. coli* RF2 to the ns70S, the sequence differences in the switch loop are unlikely to stabilize the open conformation of RF2 on the ribosome. The lack of interspecies compatibility of ResQ (and ArfA) contrasts with other, more ubiquitous ribosome rescue factors. For example, *B. subtilis* tmRNA is functional in *E. coli*, albeit with lower efficiency^25^, and human mitochondrial ArfB homolog, ICT1, is functional in *C. crescentus* and *vice versa*^23^. This low interspecies compatibility of the RF-dependent rescue factors reinforces the notion that these rescue systems emerged late in evolution.

It is crucial for the ribosome rescue factors not to intervene during normal translation, nor when translation is arrested by nascent polypeptides for regulatory purposes^32^. Our cryo-EM structure of the ResQ-RF2-ns70 reveals that ResQ uses its C-terminal region to monitor the vacancy of the ribosomal mRNA channel, a hallmark of the ribosomes stalled on the 3’ end of mRNA, and thereby discern non-stop translation complexes from those of ongoing translation^41^. Consistently, we have shown that neither ResQ, nor *trans*-translation, can resolve the MifM-programmed elongation arrest complexes (**Supplementary Fig. 1**). In this case, the mRNA channel should remain occupied by the *mifM* mRNA. Also, the cryo-EM structures of the MifM-stalled ribosomes reveal altered conformations of 23S rRNA residues near the peptidyl transferase center (PTC) such that they block accommodation of aminoacyl-tRNAs or mimics thereof into the A-site^52^ explaining why MifM stalling is refractory to the ResQ-RF2 and the SmpB-SsrA actions, too.

We have shown that *resQ* is transcribed as a non-stop mRNA because of the terminator sequence within the coding region. Thus, *resQ* itself is subject to a futile type of translation, producing a non-stop polypeptide that is extended by the SsrA tag sequence and rapidly eliminated by proteolysis. ResQ accumulates only when *trans*-translation is impaired, and in the form of peptidyl-tRNA. However, once some free ResQ product is generated by spontaneous hydrolysis of the ResQ peptidyl-tRNA, or by the presence of residual ResQ in the cell, it actively liberates *in trans* the ResQ peptide in a self-perpetuating manner. We have shown clearly using biochemical assays that a liberated form of the non-stop ResQ peptide (residues 1-62) is active in mediating the RF2-dependent peptidyl-tRNA hydrolysis.

Because ResQ is not effectively produced in *trans*-translation proficient cells, but induced strikingly upon dysfunction of *trans*-translation, it is likely to represent a secondary, back-up rescue system that compensates for defects in *trans*-translation. This scenario reinforces the notion that the proteolytic function characteristically associated with the SmpB-SsrA system is not essential for growth, as reported previously^13,14^. Indeed, the GFP-ns non-stop product accumulates in the *ssrA*-deleted cell (**Fig. 4a**), indicating that the products of the ResQ system are not necessarily toxic. Thus, the growth-essential roles of *trans*-translation and ResQ are in their ribosome recycling functions, rather than in the tagging-proteolysis that the ResQ system lacks. Cell viability would require a sufficient pool of the uncompromised ribosomes, which maintains the translation capacity of the cell. Whereas the two pathways share the essential function required for growth-supporting ribosome rescue, the proteolytic functions of the *trans*-translation and that of the RqcH tail-adding system could become more important under more severe stress conditions^49^.

The regulatory scheme of ResQ expression is strikingly similar to that elucidated for ArfA regulation in *E. coli* ^17,18^; the *arfA* mRNA is also subject to RNase III-dependent cleavage and/or transcription termination such that ArfA only accumulates when *trans*-translation is defective. It is noteworthy that the two evolutionarily unrelated rescue factors employ a similar scheme of regulation. Convergent acquisition of such regulatory mechanisms may be more common for factors that have evolved recently and which have functions related to the firmly established and relatively rigid constituent of the cell, such as the ribosome and translation factors.

In summary, our study reveals that bacteria, both Gram-negative and Gram-positive, have RF-dependent mechanisms of ribosome rescue that allow for the stop-codon independent liberation of the polypeptide from the ribosome on the non-stop mRNA (**Fig. 6a**). However, the crucial adapter proteins, ArfA, ArfT and ResQ, are unrelated in amino acid sequence (**Supplementary Fig. 2**). The modes of their interactions with the catalytic RF partner(s) are also divergent, indicative of the tailored nature of their evolution in different species. It remains possible that many more factors in this category exist in different organisms. As the present study suggests that sequence similarity alone cannot be used to identify additional factors that exist in the domains of life, we need better strategies to address this question. Further studies on the generality and diversity of independently evolved rescue factors and their manner of interaction with the ribosome and translation factor would provide invaluable insights into regulatory mechanisms of translation processes in the cell.

## Methods

### Bacterial Strains and Plasmids

*B. subtilis* and *E. coli* strains, plasmids, DNA oligonucleotides used in this study are listed in Supplementary Tables 2, 3, 4 and 5, respectively. The *B. subtilis* strains were derivatives of PY79 (wild-type; ref.^53^) and constructed by transformation that involves homologous recombination with plasmids listed in Supplementary Table 6. These plasmids carried an engineered *B. subtilis* gene to be integrated, which was flanked by sequences from the integration target loci, and were constructed by standard cloning methods including PCR, PrimeSTAR mutagenesis (Takara), and Gibson assembly^54^. The plasmid pCH747 was constructed by cloning an SphI-SpeI fragment of pCH735 into pyqjG21. Plasmids pCH735 and pyqjG21 were constructed as described^34^. Successful integration of a gene into the chromosome was accomplished by double crossing-over at the target loci. The resulting recombinant clones were checked for their antibiotic resistance markers, including the absence of those originally present on the plasmid backbone, and inactivation of the *amyE*, *lacA* or *thrC* target locus. The marker-less deletion mutants of *smpB*, *yesZ* and *lacA* were constructed by excising the drug resistance gene cassette by the Cre-loxP system as described previously^27^ with some modification as follows. The *B. subtilis* strains were transformed with pMK2, a pLOSS*-based Ts plasmid harboring *cre*. The resulting strain was grown at 37°C overnight in LB agar medium supplemented with 1 mM IPTG (isopropyl β-D-1-thiogalactopyranoside) and 100 μg/ml spectinomycin to excise the drug marker flanked by *loxP*^27^. The strain was then grown at 37°C overnight to drop off pMK2 on LB agar medium without spectinomycin. The absence of the drug resistance confirmed the absence of plasmid pMK2. The *B. subtilis* strain KFB792 was constructed by transformation of PY79 with a DNA fragment prepared by Gibson assembly with three PCR fragments, one of which was amplified from pCH1142 using a pair of primers SP89/SP90, and the other two of which were amplified from PY79 genomic DNA using pairs of primers SP91/SP92 and SP93/SP94, respectively.

### Growth conditions and general procedures

For Western blotting in Fig. 2c, *B. subtilis* cells were cultured at 37°C in LB medium with or without 0.5 % xylose until OD_600_ reached ~0.5. Bacterial culture (1 mL) was treated with 5 % trichloroacetic acid (TCA), and precipitates formed were washed with 0.75 mL of 1 M Tris-HCl (pH 8.0) and resuspended in 50 μl of buffer L (33 mM Tris-HCl, 1 mM EDTA, pH8.0) containing 1 mg/mL lysozyme, followed by incubation at 37°C for 10 min. Proteins were then solubilized with an equal volume of 2xSDS-loading buffer containing 5 mM dithiothreitol (DTT) with incubation at 65°C for 5 min and subjected to SDS-PAGE and immunoblotting.

### Synthetic lethal screening using the BKE strain library

To isolate *B. subtilis* mutants whose viability depends on *trans*-translation, we used the BKE library (a collection of single-gene knockout mutants covering the 3,968 non-essential genes, which had been disrupted by replacement with the erythromycin resistance marker) as the source of gene knockouts. We pooled the BKE strains and prepared a genomic DNA mixture using Wizard genome DNA purification kit (Promega). We used this DNA preparation to transform an *smpB*-deleted strain of *B. subtilis* that harbored a rescue plasmid carrying *smpB*^+^ and *lacZ*^+^ (pNAB1286), which was constructed from pLOSS* with a temperature sensitive (Ts) replication system. Transformant mixture was then incubated at 50°C overnight to segregate out the Ts plasmid from bacteria that did not need the rescue plasmid, followed by further growth at 37°C overnight and plating on LB agar containing 40 μg/mL X-Gal (5-bromo-4-chloro-3-indolyl β-D-galactopyranoside), 1 mM IPTG, 12.5 μg/mL lincomycin and 1 μg/mL erythromycin. We picked up blue (*lacZ*^+^) colonies to obtain transformants that retained the *smpB*^+^-*lacZ*^+^ rescue plasmid even after the high-temperature incubation and prepared chromosomal DNA from them. Genes that had been disrupted by the erythromycin-resistance marker were determined by PCR amplification and DNA sequencing of the mutant-specific barcode sequence, using appropriate primers^27^.

### CRISPR interference

The CRISPRi was performed as described previously^55^ with some modification. A gene encoding dCas9 under the xylose-inducible promoter was integrated into the *lacA* site on the chromosome of a *resQ*-deleted *B. subtilis* strain, into which an *ssrA*-targeted or an *smpB*-targeted sgRNA gene under a constitutive promoter was further integrated at the *amyE* site. The guide sequences for CRISPRi were designed on the basis of information provided by the previous study^55^ as well as CHOPCHOP, a web tool for the CRISPR/Cas9 experiments^56,57^. The target gene knock-down was induced by addition of 1% xylose.

### Protein purification

Hexahistidine-tagged proteins (*B. subtilis* ResQ62-His_6_, RF1-His_6_, RF2-His_6_, *E. coli* His_6_-ArfA60 and their derivatives) were expressed in *E. coli* strain BL21(DE3) from the pET28b-based plasmid. *E. coli* cells were grown in LB-kanamycin (25 μg/mL) medium. At a mid-log phase, IPTG (final concentration, 1 mM) was added, and cells were grown for an additional 3 hours to express and accumulate the target protein. Cells were then harvested, washed with ice-cold 50 mM HEPES-NaOH (pH7.6) buffer and stored at −80°C. They were suspended in binding buffer (50 mM HEPES-KOH pH7.6, 5 mM imidazole, 300 mM NaCl, 1 mg/mL Pefabloc) and disrupted by passing through a microfluidizer LV1 (Microfluidics) at 16,000 psi three times. After removal of debris by centrifugation (4°C, 15,000 rpm for 15 min), Ni-NTA agarose was added to the sample, which was then incubated at 4°C for 1 hour. The Ni-NTA agarose was loaded on a spin column and washed seven times with wash buffer (50 mM HEPES-KOH pH7.6, 20 mM imidazole, 0.1% Triton-X). Protein was eluted with the elution buffer (20 mM HEPES-KOH pH7.6, 300 mM NaCl, 300 mM imidazole). Purified release factors were dialyzed against dialysis buffer A (50 mM HEPES-KOH pH7.6, 100 mM potassium acetate, 1mM DTT, 30% glycerol). Purified ResQ62-His_6_ and His_6_-ArfA(2-60) were dialyzed against dialysis buffer B (50 mM HEPES-KOH pH7.6, 100 mM potassium acetate, 1 mM DTT, 300 mM NaCl, 30% glycerol).

For the structural analysis, the wildtype *B. subtilis* RF2 and variant RF2-GGP protein were expressed from pET11a vectors incorporating a C-terminal hexa-histidine tag (His_6_) for purification and detection purposes. The inactive RF2-GGP mutant was generated by site-directed mutagenesis. The wildtype RF2 and RF2-GGP proteins were over-expressed in *E. coli* BL21 (DE3) at 37 °C for 1.5 h after induction with 1 mM IPTG. Cells were collected and the pellet was re-suspended in lysis buffer (50 mM NaH_2_PO_4·_, 300 mM NaCl, 5 mM imidazole, pH 7.5). Lysis was performed using a microfluidizer (Microfluidics M-110L) by passing cells three times (at 18,000 psi). The cell debris was removed upon centrifugation and the proteins were purified from the supernatant by His-tag affinity chromatography using Ni-NTA agarose beads (Clontech). The bound proteins were washed with lysis buffer containing 10 mM imidazole and then eluted with lysis buffer containing 250 mM imidazole. The proteins RF2, RF2-GGP and ResQ62-His_6_ were purified by size-exclusion chromatography using HiLoad 16/600 Superdex 75 (GE Life Sciences) in gel filtration buffer (50 mM HEPES, pH 7.4, 50 mM KCl, 100 mM NaCl, 2% glycerol, 5 mM β-mercaptoethanol). The proteins were concentrated using Amicon Ultracel-30 Centrifugal Filter Units (Merck Millipore) for wild-type RF2 and RF2-GGP and Ultracel-3 for ResQ62-His_6_ hereafter referred to as ResQ.

### *In vitro* translation using PURE system

The *E. coli*-based coupled transcription-translation system with purified components (PUREfrex 1.0; GeneFrontier) was used for in vitro translation as described previously^29,30,58^ with some modifications. To maximize transcription, we added 2.5 U/μL of T7 RNA polymerase (Takara) further to the reaction mixture. Whereas the original reaction mixture, referred to as *Ec* PURE system, contained the *E. coli* ribosome, we also used *Bs* hybrid PURE system containing the *B. subtilis* ribosomes at a final concentration of 1 μM. Unless otherwise noted, we omitted RF1, RF2 and RF3 from the reaction. However, we included purified release factor or its derivatives derived either from *B. subtilis* or *E. coli* at a final concentration of 1 μM as indicated in each experiment. *E. coli* RFs were purchased from GeneFrontier. Purified ResQ or ArfA was added to the final concentration of 1 μM when indicated. The reaction was primed with an appropriate DNA fragment prepared by PCR (Supplementary Table 7) and allowed to continue at 37°C for 20 min. Samples were then mixed with the same volume of 2xSDS-PAGE loading buffer. When indicated, they were further treated with 0.2 mg/mL RNaseA (Promega) at 37°C for 15 min before electrophoresis. Samples for SDS-PAGE were heated at 65°C for 5 min and separated by 10% wide range gel (Nacalai Tesque). Translation products were detected by Western blotting using anti-GFP (A-6455; Thermo) or anti-DYKDDDDK (anti-FLAG tag; Wako) as described previously^58^.

### Generation of ResQ-RF2-ns70S complex

Generation of the ResQ-RF2-ns70S complex was similar to that previously described for the ArfA-RF2-70S complex^38^. Briefly, the truncated nlpD template containing an N-terminal His_6_ and HA-tag was first amplified from pET21b-r1nlpD using T7-promotor and 133-159 nlpD reverse primer (**Supplementary Table 6**). Following PCR purification via spin column (Qiagen), *in vitro* translation was performed using the PURExpress *In vitro* Protein Synthesis kit (NEB 6800) and the non-stop ribosome complex was purified as previously described in ref.^38^. The purified non-stop ribosome complex was then incubated together with a 10x excess of ResQ and RF2-GGP mutant for 5 min at 37°C before being applied to cryo-EM grids.

### Cryo-EM and single particle reconstruction

Three microliters (4.5 OD_260nm_ per mL) of ResQ-RF2-ns70S complex was applied to 2nm pre-coated Quantifoil R3/3 holey carbon supported grids and vitrified using the Vitrobot Mark IV (FEI, Holland). Data collection was performed using EM-TOOLS (TVIPS GmbH) on a Titan Krios transmission electron microscope equipped with a Falcon III direct electron detector (FEI, Holland) at 300 keV at a pixel size of 1.065 Å and a defocus range of 0.4–2.2 µm. Forty frames (dose per frame of 2 e^−^ Å^−2^) were aligned using Motion Correction software^59^. Power-spectra and defocus values were determined using the GCTF software^60^. Micrographs showing thon rings beyond 3.2 Å were manually inspected for good areas and automatic particle picking was performed using the Gautomatch software (http://www.mrclmb.cam.ac.uk/kzhang/). Single particles were then imported and processed in Relion 3^61^. 514,119 particles were first subjected to 2D classification (60 classes for 100 rounds) and 389,381 particles showing ribosome-like features were then selected for 3D refinement using an *E. coli* 70S ribosome as a reference structure (**Supplementary Fig. 4a**). 3D classification was then performed, resulting in 317,095 particles containing P-site tRNA and RF2 (**Supplementary Fig. 4b**) that were further selected for focus sorting on the RF2 (**Supplementary Fig. 4c**). Focus sorting yielded a major population of 154,405 particles containing stoichiometric amounts of ResQ, P-site tRNA, RF2, which after a final round of 3D refinement produced a final cryo-EM reconstruction with an average resolution of 3.15 Å according to FSC_0.143_ criterion (**Supplementary Fig. 4d**). The final cryo-EM maps were sharpened by dividing the maps by the modulation transfer function of the detector and applying an automatically determined negative B factor in Relion 3^61^. The final cryo-EM map was also filtered according to local resolution using SPHIRE^62^.

### Molecular modeling of the ResQ-RF2-ns70S complex

The molecular model for the ribosomal proteins and rRNA core was based on the molecular model from the recent cryo-EM reconstructions of the *E. coli* 70S ribosome (PDB ID 6H4N^63^ and 5MGP^38^. The models were rigid body fitted into the cryo-EM density map using UCSF Chimera followed by refinement in Coot^64^. Proteins L1, L10, L11 protein and the L7/L12 stalk were not included in the final model due to the poor quality of density in the final cryo-EM map. For *B. subtilis* RF2, a homology model was generated using HHPred^65^ based on an *E. coli* RF2 template (PDB ID 5MGP^38^). Domains 1 and 3 of RF2 were less well-resolved due to high flexibility (**Supplementary Fig. 4e**) and therefore only the backbone was modeled. Residues 2-55 of ResQ were built *de novo* using an HHPred model^65^ as an initial starting point to determine the placement of the central helical regions. The complete atomic model of the ResQ-RF2-ms70S complex was manually adjusted using Coot^64^ and refined with phenix real_space_refine for cryo-EM^64^ using restraints obtained by phenix secondary_structure_restraints^64^. The model refinement and statistics of the refined model were obtained using MolProbity^66^ (**Supplementary Table 1**).

### Figure preparation

Figures showing electron densities and atomic models were generated using either UCSF Chimera, UCSF ChimeraX^67^ or PyMol (Version 1.8 Schrödinger). The comparison of ResQ with mRNA (PDB ID 4V5E^47^), ArfA (PDB ID 5MGP^38^, 5MDV^36^, 5U9F^43^), RF2_closed_ and RF2_stop_ (PDB ID 4V5E^47^) was obtained by alignment of the 16S rRNAs from the respective structures using PyMol (Version 1.8 Schrödinger).

### Data availability

The cryo-EM map of the ResQ-RF2-ns70S complex is available through the EMDB code EMD-00000 and the associated molecular model is deposited in the Protein Data Bank with the entry code XXXX. The data that support the findings of this study are available from the corresponding authors on request.

## Acknowledgments

We thank Dr. Hyouta Himeno of Hirosaki University for the *B. subtilis ssrA* mutant strain, Dr. Jiří Nováček (CEITEC, Brno, Czech Republic) for help with high-resolution cryo-EM data collection, and Yuko Sumino, Saori Amano, Sakika Otani, Toru Irie, S. Rieder and C. Ungewickell for expert technical support. We are also thankful to the National BioResource Project for providing the BKE strain library. This work was supported by grants from MEXT and JSPS Grant-in-Aid for Scientific Research (Grant No. 25291006 and 16H04788 to SC, 26116008 to SC and KI, 19K16044 to KF) and the Deutsche Forschungsgemeinschaft (WI3285/4-2 to DNW).

## Author Contributions

N.S.C., K.F., and S.C. performed biochemical experiments and C.M., B.B. and D.N.W. performed structural experiments. All authors designed experiments, analyzed data and were involved in writing the manuscript.

## Additional information

Supplementary information is available for this paper at XXXX

