## Supplementary Information for "ResQ, a release factor-dependent ribosome rescue factor in the Gram-positive bacterium *Bacillus subtilis*"

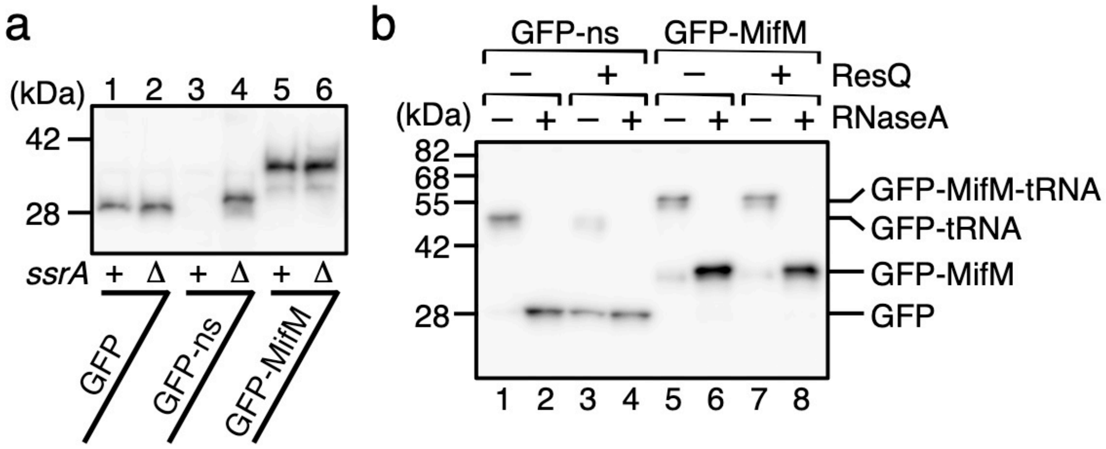

**Supplementary Fig. 1 | Nascent-chain mediated ribosome stalling is refractory to the ribosome rescue systems in *B. subtilis*.** **a**, MifM is refractory to *trans*-translation. Cellular accumulation of GFP-MifM was examined in the wild type and *ssrA*-deficient mutant strains. GFP (lanes 1, 2), GFP-ns (lanes 3, 4) and GFP-MifM (lanes 5, 6) were expressed in the wild type (odd number) and Δ*ssrA* (even number) strains of *B. subtilis*, separated by SDS-PAGE and detected with anti-GFP immunoblotting. **b**, Inability of ResQ to induce RF2 hydrolysis of the elongation-arrested MifM-tRNA. In vitro translation using *Bs* hybrid PURE system with RF2 was directed by the *gfp-ns* (lanes 1-4) or *gfp-mifM* (lanes 5-8) template in the presence (lanes 3, 4, 7, 8) or the absence (lanes 1, 2, 5, 6) of purified ResQ62-His$_6$. The translation products were analyzed by anti-GFP immunoblotting as described above.

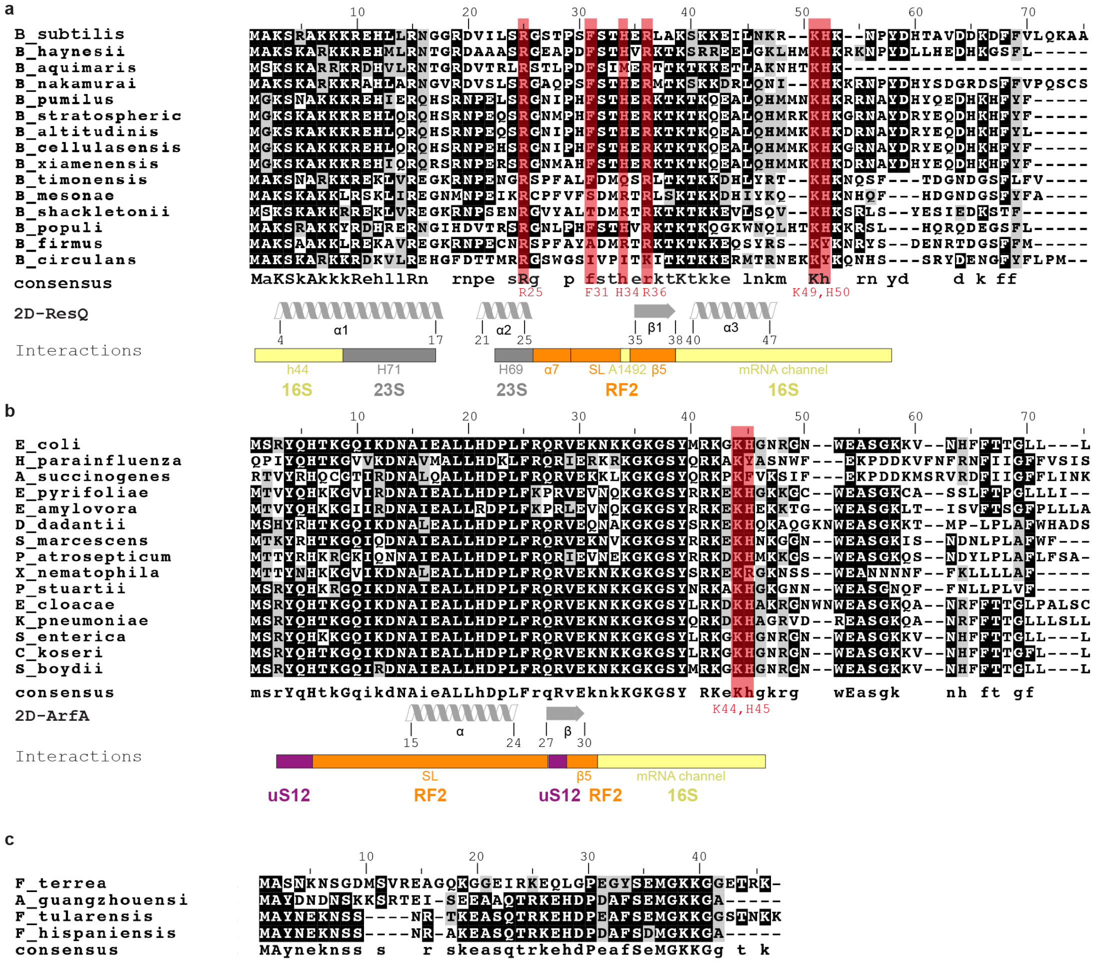

**Supplementary Fig. 2 | Multiple sequence alignments of ResQ, ArfA, and ArfT homologs.** **a-c**, Multiple sequence alignments of (**a**) ResQ, (**b**) ArfA and (**c**) ArfT generated with Clustal Omega (Madeira et al., 2019) and Boxshade with residues shaded that are identical (black) or similar (grey). Below each alignment, the consensus sequence with capital letters indicating 100% conservation and lower-case letters indicating less conservation). The known secondary structure is shown for *B. subtilis* ResQ (2D-ResQ) in (**a**) and *E. coli* ArfA (2D-ArfA) in (**b**), as well as interaction regions between (**a**) ResQ and (**b**) ArfA with the ribosome (16S rRNA, yellow, 23S rRNA, grey and uS12, violet) and RF2 (orange).

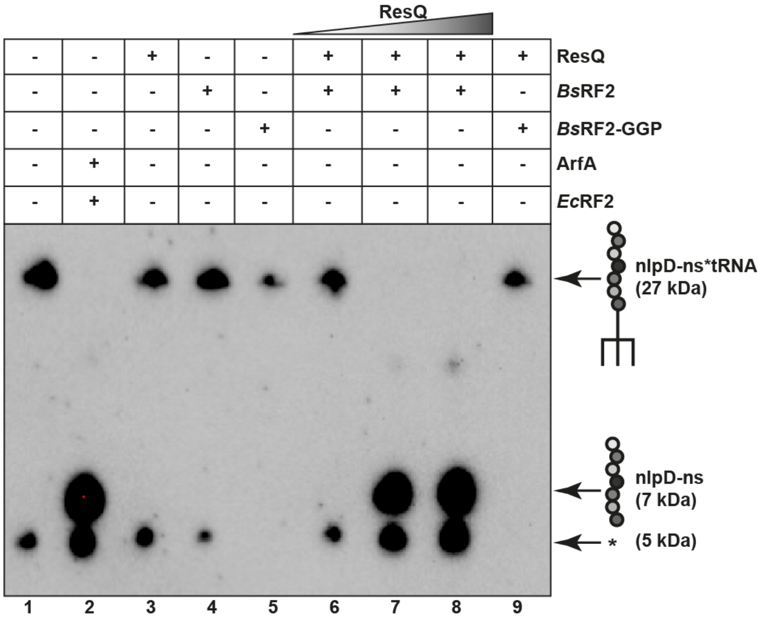

**Supplementary Fig. 3 | Recycling of non-stop ribosomes by ResQ and RF2.**

*In vitro* translation assays using the *E. coli* PURE systemΔRF123 kit (lacking all RFs) were performed with truncated non-stop *nlpD* DNA template, in the presence of ResQ, wildtype *B. subtilis* RF2 (*Bs*RF2) and the RF2-GGP mutant (*Bs*RF2-GGP) alone (lanes 3-5), or *Bs*RF2 (20 pmol) with increasing concentrations (12-100 pmol) of ResQ (lanes 6-8), or ResQ (100 pmol) in the presence of 100 pmol *Bs*RF2-GGP (lane 9). As a positive control, reactions were performed with *E. coli* ArfA and RF2 (*Ec*RF2), as described previously (lane 2) (Huter et al., 2017b). A negative control was also performed where reactions lacked all RFs and rescue factors (lane 1). Western blotting of NuPAGE gels using an antibody against the HA-tag present in the N-terminus of the NlpD peptide detected the presence of the non-stop NlpD-peptidyl-tRNA (27 kDa) and released NlpD peptide (7 kDa). The asterisk (*) indicates a mysterious band that cross-reacts with the HA-antibody, but is also present in the negative control and was therefore not examined further. As expected, the negative control (lane 1), as well as reactions performed in the presence of ResQ, *Bs*RF2 or *Bs*RF2GGP alone (lanes 3-5), show a strong band for the NlpD-peptidyl-tRNA and no evidence for the released NlpD peptide. By contrast, the positive control with ArfA and *Ec*RF2 (lane 2), as well as the reactions with *Bs*RF2 and increasing concentrations of ResQ (lanes 7 and 8) shows no NlpD-peptidyl-tRNA and only the presence of released NlpD peptide. As expected, substitution of wildtype *Bs*RF2 with the inactive *Bs*RF2-GGP mutant led to a loss the band for the released NlpD peptide and the presence of NlpD-peptidyl-tRNA was restored (lane 9).

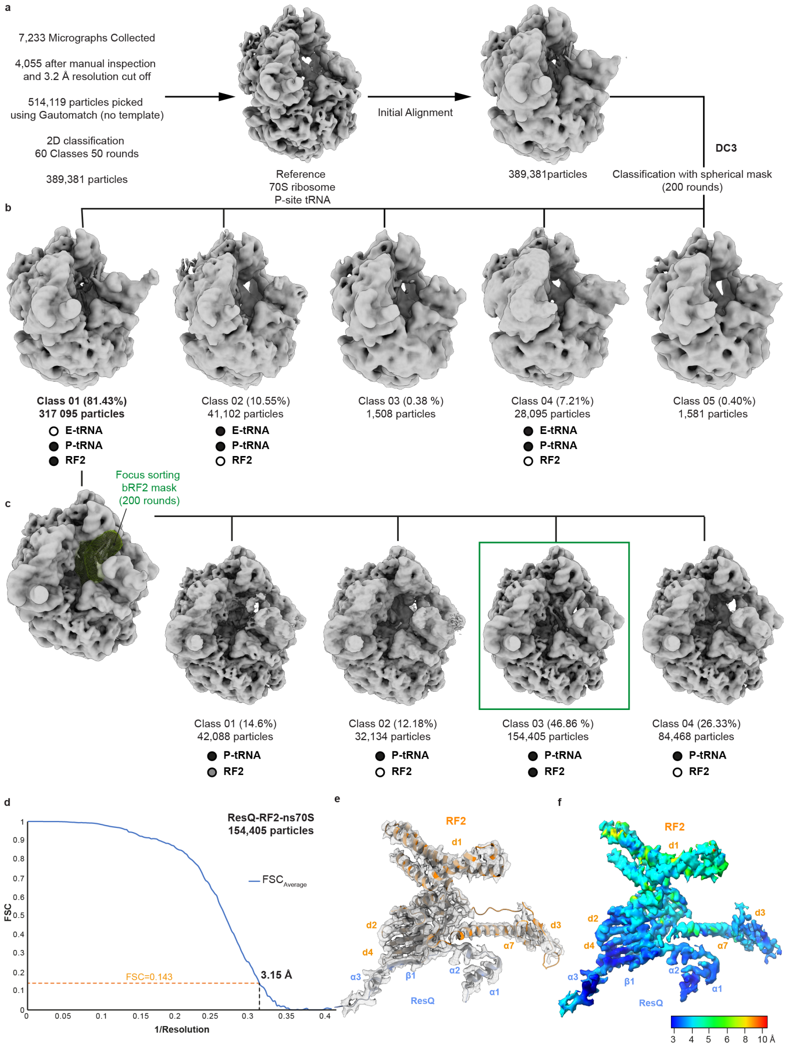

**Supplementary Fig. 4 | *In silico* sorting of the ResQ-RF2-ns70S complex**

**a**, After particle picking and extensive 2D classification, the complete dataset of 389,381 particles was initially aligned against a P-site tRNA containing *E. coli* 70S ribosome. **b**, Following 3D classification for 200 rounds in Relion, five classes were generated. The majority (317,095 particles; 81.43%) of particles were found in class 1 and contained the ResQ-RF2-ns70S complex. The second major (41,102 particles; 10.55%) class 2 contained a fully programmed ribosome, but without the presence of RF2. In addition, three minor classes 4-6 (class 3; 1,508 particles; 0.38 %, class 4; 28,095 particles; 7.21%; class 5; 1,581 particles; 0.4%) containing damaged and/or poorly aligning particles were observed. **c**, The 317,095 particles from the class 1 were further sorted using a focus sorting mask around RF2, resulting in four additional classes, of which class 3 (154,405 particles, 46.86%) contained stoichiometric occupancy of P-site tRNA, RF2 and ResQ. Class 3 was then refined to yield a (**d**) final reconstruction of the ResQ-RF2-ns70S complex with an average resolution of 3.15 Å (0.143 FSC). **e-f**, Isolated electron density for ResQ and RF2 **e**, shown as grey mesh with fitted model and (**f**) colored according to local resolution

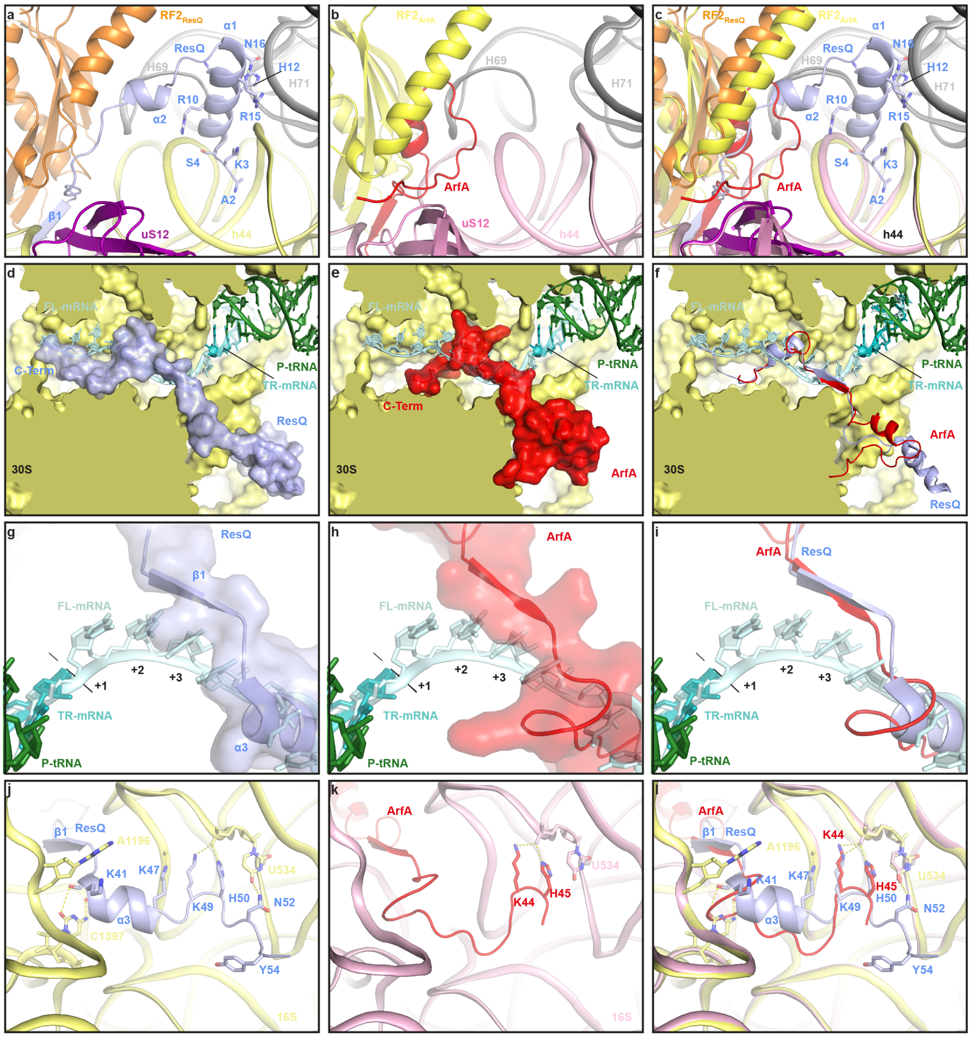

**Supplementary Fig. 5 | Interactions of ResQ and ArfA on the ribosome**. **a**, Interaction of the N-terminus of ResQ (blue) with RF2 (orange), uS12 (violet), helix 44 (h44) of the 16S rRNA (yellow) and helix 69 (H69) and H71 of the 23S rRNA (grey). **b**, Same view as (**a**), but for the ArfA-RF2-70S complex, with ArfA (red), RF2 (yellow) and 16S rRNA (pink) (Huter et al., 2017b). **c**, Overlay of (**a**) and (**b**). **d-e**, Transverse section of the 30S subunit (yellow) to reveal the mRNA channel showing a superimposition of full-length mRNA (FL-mRNA, cyan) with truncated non-stop mRNA (TR-mRNA, teal), P-site tRNA (green) and surface representations of (**d**) ResQ (blue) and (**e**) ArfA (red) (Huter et al., 2017b). **f**, Overlay of (**d**) and (**e**) with cartoon representations of ResQ (blue) and ArfA (red). **g-h**, Superimposition of full-length mRNA (FL-mRNA, cyan) with truncated non-stop mRNA (TR-mRNA, teal), P-site tRNA (green) and transparent surface representations of (**g**) ResQ (blue) and (**h**) ArfA (red) (Huter et al., 2017b). The first (+1), second (+2) and third (+3) nucleotides of the A-site codon of the FL-mRNA are indicated. **i**, Overlay of (**g**) and (**h**) with cartoon representations of ResQ (blue) and ArfA (red). **j-k**, Interaction of the C-terminus of (**j**) ResQ (blue) and (**k**) ArfA (red) (Huter et al., 2017b) with the 16S rRNA with potential hydrogen bonds indicated with yellow dashed lines. **l**, Overlay of (**j**) and (**k**).

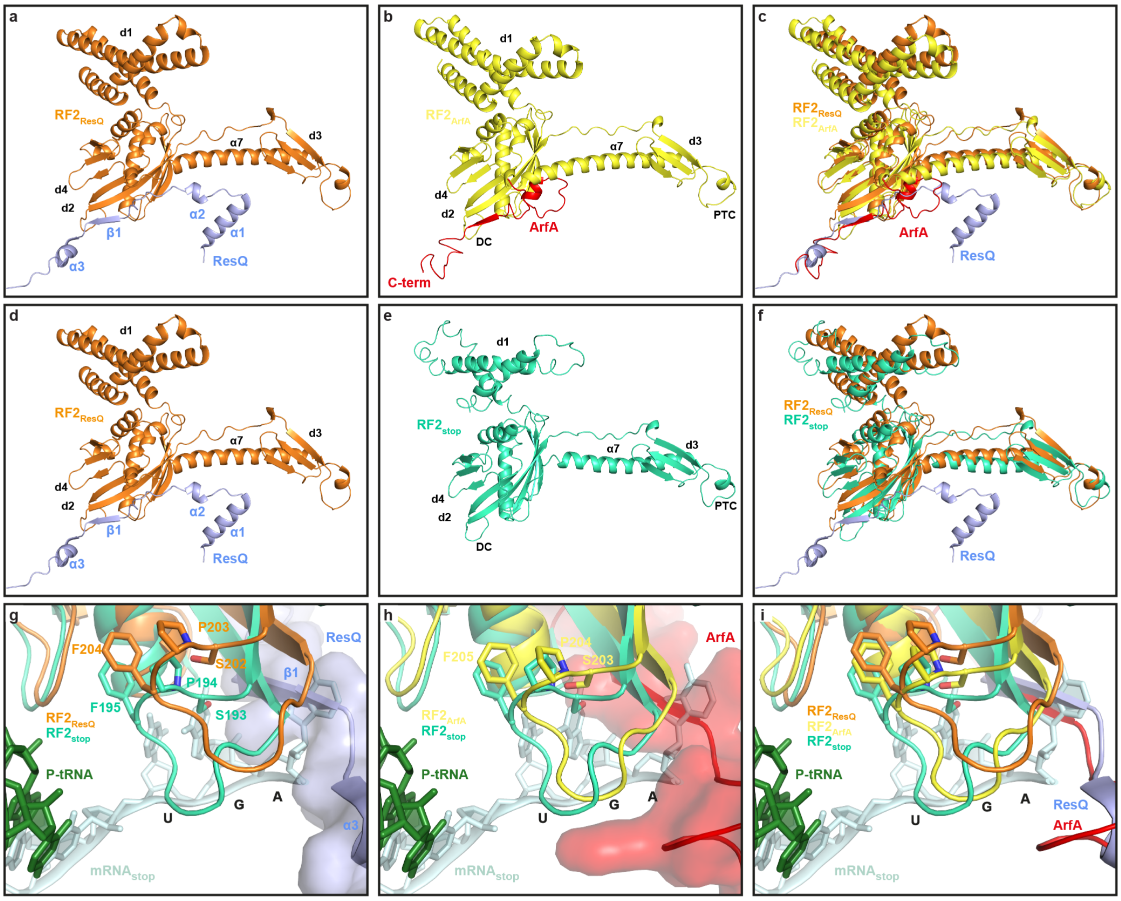

**Supplementary Fig. 6 | Interaction of ResQ and ArfA with RF2 on the ribosome**.

**a-b**, Interaction between (**a**) ResQ (blue) and RF2$_{ResQ}$ (orange) in the ResQ-RF2-ns70S complex compared with (**b**) ArfA (red) and RF2$_{ArfA}$ (yellow) in the ArfA-RF2-ribosome complex (Huter et al., 2017b). The approximate positions of the decoding center (DC) and peptidyltransferase center (PTC) on the ribosome are indicated. **c**, Superimposition of (**a**) and (**b**). **d-e**, The binding position of (**d**) RF2$_{ResQ}$ (orange) and ResQ (blue) in the ResQ-RF2-ns70S complex, compared with (**e**) RF2$_{stop}$ (lime) in a canonical termination complex (Weixlbaumer et al., 2008). The approximate positions of the decoding center (DC) and peptidyltransferase center (PTC) on the ribosome are indicated. **f**, Superimposition of (**d**) and (**e**). **g-h**, Superimposition of the SPF motif of RF2$_{stop}$ (lime) and UGA codon of the mRNA (cyan) (PDB ID 4V5E, (Weixlbaumer et al., 2008)) with (**g**) the SPF motif of RF2$_{ResQ}$ (orange) and ResQ (blue), and (**h**) with the SPF motif of RF2$_{ArfA}$ (yellow) and ArfA (red) (Huter et al., 2017b). **i**, Overlay of (**g**) and (**h**) with ResQ (blue) and ArfA (red) as cartoon representations.

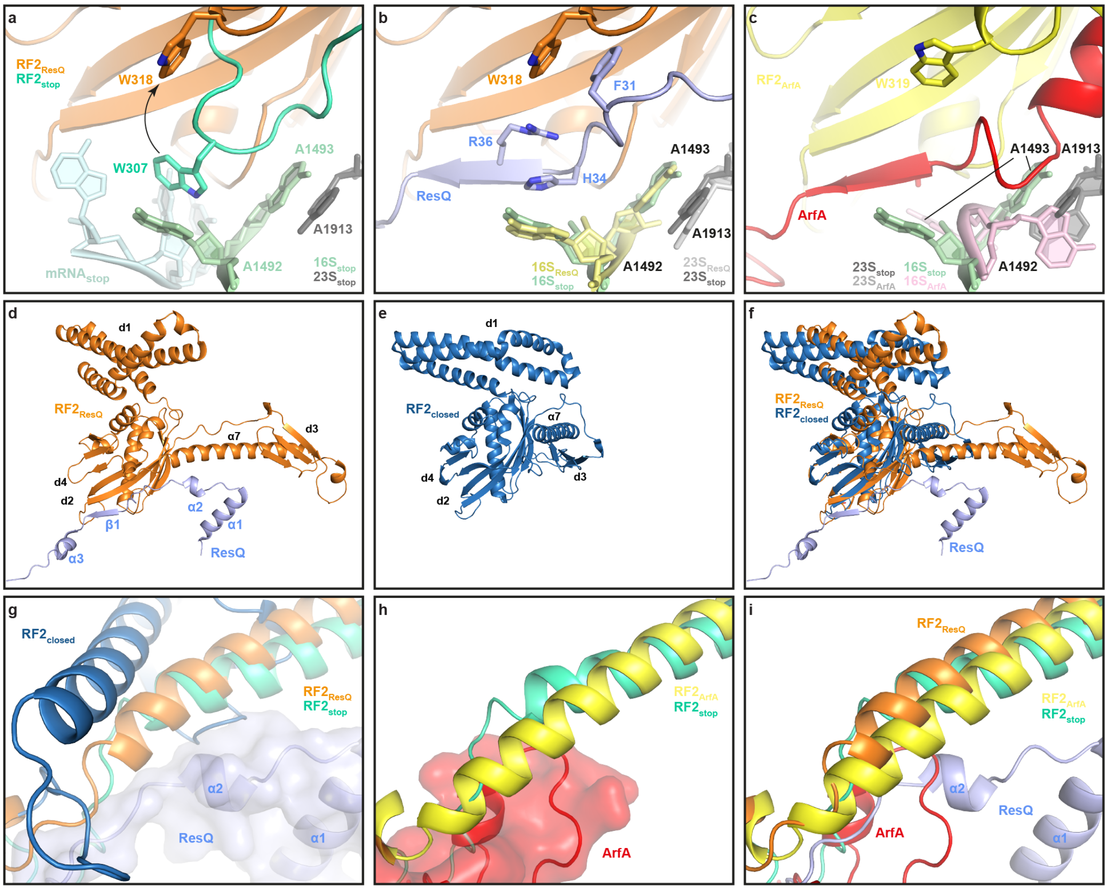

**Supplementary Fig. 7 | ResQ induces an open conformation of RF2 on the ribosome**

**a**, Interaction between Trp307 (W307, which is equivalent to *B. subtilis* Trp318 (W318)) of the switch region of *Thermus thermophilus* $RF2_{stop}$ (lime) and A1492 of the 16S rRNA (green) during decoding of the UGA stop codon of the mRNA (cyan; PDB ID 4V5E, (Weixlbaumer et al., 2008)). W318 in the switch loop of RF2 ($RF2_{ResQ}$, orange) observed upon ResQ binding is superimposed and arrowed. **b**, Same view as in (**a**), showing the conformation of the switch loop of $RF2_{ResQ}$ (orange) and A1492/A1493 (pale yellow) when ResQ (blue) is present. **c**, Same view as in (**a**) and (**b**), showing the conformation of A1492/A1493 in comparison to (**a**) and the switch loop in the presence of ArfA (red, PDB ID 5MVGP, (Huter et al., 2017b). (**d**) Open conformation observed for $RF2_{ResQ}$ (orange) when in complex with ResQ on the ribosome, compared with (**e**) the closed conformation of $RF2_{closed}$ (dark blue, PDB ID 1GQE, (Vestergaard et al., 2001)) when not bound to the ribosome. **f**, Superimposition of (**d**) and (**e**). **g**, Superimposition of the conformation of helix α7 of RF2 from the crystal structure of the closed form of $RF2_{closed}$ with $RF2_{Stop}$ (lime) and $RF2_{ResQ}$ (orange), with ResQ (blue) shown for reference. **h-i**, Same view as in (**g**) showing the conformation of helix □7 of $RF2_{stop}$ superimposed with (**h**) $RF2_{ArfA}$, and ArfA (red) and (**i**) including $RF2_{ResQ}$ (orange) and ResQ (blue).

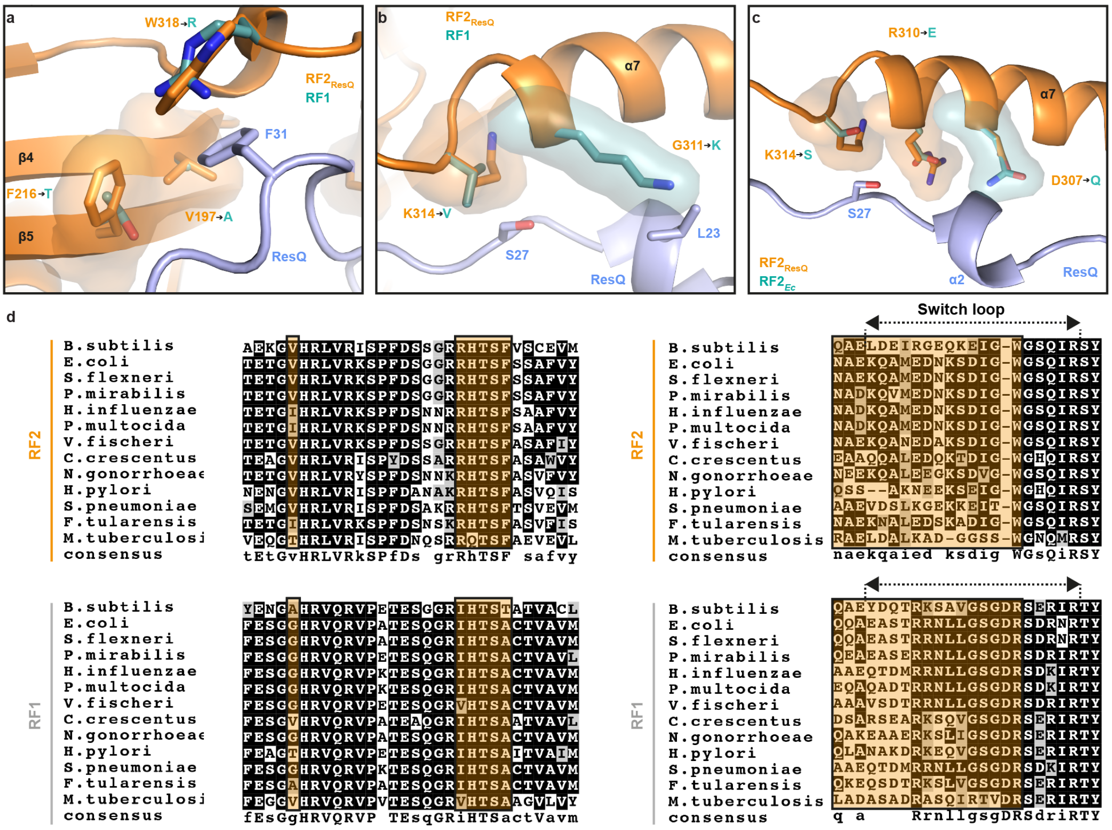

**Supplementary Fig. 8 | Basis for the species-specificity of ResQ**

(**a-b**) Potential species-specific interactions between ResQ (blue) and *B. subtilis* RF2 (orange) compared with a homology model for *B. subtilis* RF1 (green) aligned to the ResQ-RF2-ns70S. **a**, The ResQ interface with β4 and β5 strands of *B. subtilis* RF2 (orange) consists of hydrophobic residues Val197 and Phe216, which are substituted by Ala and Thr, respectively in *B. subtilis* RF1. **b**, The ResQ interface with the switch loop and helix α7 of *B. subtilis* RF2 (orange) consists of multiple residues that are distinct *B. subtilis* RF2 and RF1, for example, G311, K314 and W318 (seen in (**a**)) of RF2 that are substituted with Lys, Val and Arg, respectively, in *B. subtilis* RF1. **c**, Potential sequence differences between *B. subtilis* RF2 (orange) and *E. coli* RF2 (green) that could come within close proximity of ResQ and could explain the species-specific activity of ResQ. **d**, Sequence alignments of RF1 and RF2 for the corresponding regions shown in (**a**)-(**c**).

### Supplementary Tables

**Supplementary Table 1 Data collection, refinement and validation statistics**

| | **ResQ-RF2-ns70S** |
|---|---|
| **Data collection** | |
| Particles | 154,405 |
| Pixel size (Å) | 1.065 |
| Defocus range ($\mu$m) | 0.4-2.2 |
| Voltage (keV) | 300 |
| Electron dose ($e^-$/Å$^2$) | 20 |
| **Model composition** | |
| Protein resitdues | 6100 |
| RNA nucleotides | 4640 |
| Hydrogens | 0 |
| **Refinement** | |
| Resolution (Å) | 3.15 |
| Map sharpening B factor (Å$^2$) | -123.33 |
| CC map/model | 0.85 |
| **Validation: proteins** | |
| Poor rotamers (%) | 1.80 |
| Ramachandran outliers (%) | 0.42 |
| Bad backbone bonds (%) | 0.00 |
| Bad backbone angels (%) | 0.06 |
| **Validation: RNA** | |
| Correct sugar puckers (%) | 99.15 |
| Good backbone conformations (%) | 79.74 |
| Bad bonds (%) | 0.00 |
| Bad angels (%) | 0.02 |
| **Scores** | |
| MolProbity | 1.99 (76$^{th}$ percentile) |
| Clash score, all atoms | 5.64 (92$^{nd}$ percentile) |

### Supplementary Table 2. *E. coli* strains

| Name | Description | Reference |
|---|---|---|
| BL21(DE3) | $F^-$, *ompT*, *hsdS*$_B$($r_B^-$ $m_B^-$), *gal*(*λcI 857, ind1, Sam7, nin5, lacUV5-T7gene1*), *dcm*(DE3) | Promega |
| NAE970 | BL21(DE3)/pNAR913 (*Bs_prfA-his*$_6$) | This study |
| NAE972 | BL21(DE3)/pNAR915 (*Bs_prfB-his*$_6$) | This study |
| NAE973 | BL21(DE3)/pNAR916 (*resQ62-his*$_6$) | This study |
| NAE982 | BL21(DE3)/pNAR917 (*his*$_6$*-arfA(2-60)*) | This study |
| NAE1003 | BL21(DE3)/pCH2307 (*Bs_prfB(GAQ)-his*$_6$) | This study |
| NAE1017 | BL21(DE3)/pNAR939 (*Bs_pfrB(SPT)-his*$_6$) | This study |

### Supplementary Table 3. *B. subtilis* strains and construction

| Name | Description | Reference | Construction | |
|---|---|---|---|---|
| | | | Host | DNA |
| PY79 | *WT* | ref. (1) | | |
| BKE23540 | *ΔresQ::erm* | ref. (2) | | |
| BKE33600 | *ΔsmpB::erm* | ref. (2) | | |
| D1 | *ΔssrA::cat* | ref. (3) | | |
| TSB2 | *ΔssrA::cat::tet* | This study | SCB2582 | pCm::Tc |
| NAB1196 | *thrC::*$P_{xylA}$ *GFPΩerm* | This study | PY79 | pNAR778 |
| NAB1198 | *thrC::*$P_{xylA}$ *GFP-nsΩerm* | This study | PY79 | pNAR780 |
| NAB1200 | *ΔssrA::cat, thrC::*$P_{xylA}$ *GFPΩerm* | This study | SCB2582 | pNAR778 |
| NAB1202 | *ΔssrA::cat, thrC::*$P_{xylA}$ *GFP-nsΩerm* | This study | SCB2582 | pNAR780 |
| NAB1280 | *ΔsmpB::erm* | This study | PY79 | BKE33600 |
| NAB1281 | *ΔyesZ::loxP, ΔlacA::loxP, ΔsmpB::erm* | This study | KFB793 | NAB1233 |
| NAB1282 | *ΔsmpB::loxP* | This study | NAB1280 | pMK2 |
| NAB1283 | *ΔyesZ::loxP, ΔlacA::loxP, ΔsmpB::loxP* | This study | NAB1281 | pMK2 |
| NAB1286 | *ΔyesZ::loxP, ΔlacA::loxP, ΔsmpB::loxP / pNAR813* | This study | NAB1283 | pNAR813 |
| NAB1298 | *ΔresQ::erm* | This study | PY79 | BKE23540 |
| NAB1346 | *ΔresQ::kan* | This study | PY79 | pNAR903 |
| SCB851 | *amyE::*$P_{mifM}$ *rbsm1-GFP-mifM35-flag-yidC2'-lacZΩcat* | ref. (4) | | |
| SCB2582 | *ΔssrA::cat* | This study | PY79 | D1 |
| SCB4122 | *amyE::*$P_{mifM}$ *rbsm1 GFP-resQ62-FLAGΩcat* | This study | PY79 | pCH2240 |
| SCB4126 | *ΔsmpB::erm, amyE::*$P_{mifM}$ *rbsm1 GFP-resQ62-FLAGΩcat* | This study | NAB1280 | pCH2240 |
| SCB4141 | *amyE::*$P_{mifM}$ *rbsm1-GFP-resQ-FLAGΩcat* | This study | PY79 | pCH2239 |
| SCB4142 | *amyE::*$P_{mifM}$ *rbsm1-GFP-resQ(no_term)-FLAGΩcat* | This study | PY79 | pCH2241 |
| SCB4144 | *ΔsmpB::erm, amyE::*$P_{mifM}$ *rbsm1-GFP-resQ-FLAGΩcat* | This study | NAB1280 | pCH2239 |
| SCB4145 | *ΔsmpB::erm, amyE::*$P_{mifM}$ *rbsm1-GFP-resQ(no_term)-FLAGΩcat* | This study | NAB1280 | pCH2241 |
| SCB4153 | *lacA::*$P_{xylA}$ *dCas9Ωerm, amyE::*$P_{veg}$ *sgRNA-smpBΩcat* | This study | KFB946 | pCH2264 |
| SCB4191 | *lacA::*$P_{xylA}$ *dCas9Ωerm, ΔresQ::kanR* | This study | KFB946 | NAB1346 |
| SCB4194 | *amyE::*$P_{veg}$ *sgRNA-smpBΩcat, lacA::*$P_{xylA}$ *dCas9Ωerm, ΔresQ::kanR* | This study | SCB4191 | pCH2264 |
| SCB4199 | *amyE::*$P_{veg}$ *sgRNA-ssrAΩcat, lacA::*$P_{xylA}$ *dCas9Ωerm, ΔresQ::kanR* | This study | SCB4191 | pCH2302 |
| SCB4205 | *lacA::*$P_{xylA}$ *dCas9Ωerm, amyE::*$P_{veg}$ *sgRNA-ssrAΩcat* | This study | KFB946 | pCH2302 |

| | | | | |
|---|---|---|---|---|
| SCB4215 | *amyE::P*$_{veg}$ *sgRNA-smpBΩcat, lacA::P*$_{xylA}$ *dCas9Ωerm, ΔresQ::kanR, thrC::PresQ resQΩspc* | This study | SCB4194 | pCH2293 |
| SCB4217 | *amyE::P*$_{veg}$ *sgRNA-ssrAΩcat, lacA::P*$_{xylA}$ *dCas9Ωerm, ΔresQ::kanR, thrC::PresQ resQΩspc* | This study | SCB4199 | pCH2293 |
| SCB4218 | *amyE::P*$_{mifM}$ *rbsm1-GFP-mifM35-flag-yidC2'-lacZΩcat, ssrA::cat::tet* | This study | SCB851 | TSB2 |
| KFB792 | *ΔlacA::loxP-kanR* | This study | PY79 | DNA fragment |
| KFB793 | *ΔyesZ::loxP, ΔlacA::loxP* | This study | KYB112 | KFB792, pMK2 |
| KFB946 | *lacA::P*$_{xylA}$ *dCas9Ωerm* | This study | PY79 | pJMP1 |
| KFB948 | *ΔyesZ::loxP, lacA::Pxyl-dCas9Ωerm* | This study | KYB112 | pJMP1 |
| KYB112 | *ΔyesZ::loxP* | This study | PY79 | pKY13, pMK2 |

### Supplementary Table 4. Plasmids

| Name | Description | Source |
|---|---|---|
| pCm::Tc | *Cm::Tc* | ref. (5) |
| pDG1664 | *thrC::erm* integration vector | ref. (6) |
| pET28b | vector | Novagen |
| pLOSS* | Ts vector | ref. (7) |
| pyqjG21 | *amyE::P*$_{mifM}$ *mifM-yidC2-gfp* | ref. (4) |
| pJMP1 | *lacA::Pxyl dCas9Ωerm* | ref. (8) |
| pNAR756 | *P*$_{xylA}$ *yidC2* | This study |
| pNAR758 | *P*$_{xylA}$ *yidC2-non_stop* | This study |
| pNAR778 | *thrC::P*$_{xylA}$ *GFPΩerm* | This study |
| pNAR780 | *thrC::P*$_{xylA}$ *GFP-nsΩerm* | This study |
| pNAR809 | *P*$_{spac}$ *smpB* | This study |
| pNAR813 | *P*$_{spac}$ *smpB-FLAG* | This study |
| pNAR869 | *resQ-his*$_6$ | This study |
| pNAR879 | *amyE::P*$_{mifM}$ *rbsm1-GFP-resQΩcat* | This study |
| pNAR901 | upstream region of *resQ* | This study |
| pNAR903 | *ΔyqkK::kan* | This study |
| pNAR913 | *Bs_prfA-his*$_6$ | This study |
| pNAR915 | *Bs_prfB-his*$_6$ | This study |
| pNAR916 | *resQ62-his*$_6$ | This study |
| pNAR917 | *his*$_6$*-arfA(2-60)* | This study |
| pNAR939 | *Bs_pfrB(SPT)* | This study |
| pCH735 | *amyE::P*$_{mifM}$ *mifM-lacZΩcat* | ref. (4) |
| pCH747 | *amyE::P*$_{mifM}$ *mifM-gfpΩcat* | This study |
| pCH805 | *amyE::P*$_{mifM}$ *GFP-mifM35-yidC2'-lacZΩcat* | ref. (4) |

| | | |
|---|---|---|
| pCH913 | *amyE::*$P_{mifM}$ *rbsm1-GFP-mifM35-yidC2'-lacZΩcat* | ref. (4) |
| pCH1141 | *thrC::*$P_{xylA}$*Ωerm* | ref. (9) |
| pCH1142 | *kanΩspc* | ref. (10) |
| pCH2238 | *his*$_6$*-Bs_prfB* | This study |
| pCH2239 | *amyE::*$P_{mifM}$ *rbsm1 GFP-resQ-FLAGΩcat* | This study |
| pCH2240 | *amyE::*$P_{mifM}$ *rbsm1 GFP-resQ62-FLAGΩcat* | This study |
| pCH2241 | *amyE::*$P_{mifM}$ *rbsm1 GFP-resQ(no_term)-FLAGΩcat* | This study |
| pCH2264 | *amyE::*$P_{veg}$ *sgRNA-smpBΩcat* | This study |
| pCH2293 | *thrC::PresQ resQΩspc* | This study |
| pCH2302 | *amyE::*$P_{veg}$ *sgRNA-ssrAΩcat* | This study |
| pCH2307 | *Bs_prfB(GAQ)-his*$_6$ | This study |
| pKIG855 | *amyE::*$P_{veg}$ *sgRNA-rfpΩcat* | This study |
| pMK2 | *Pspac cre* | This study |
| pKY13 | *ΔyesZ::loxP-kan-loxP* | This study |
| pNR1 | *amyE::*$P_{xylA}$ *rbsm1 mifM-GFPΩcat* | This study |

### Supplementary Table 5. Primers

| Name | Sequence |
|---|---|
| SP1 | TATTTTTAAGGGGGAAATCACATAAAAAGGAGGAGAACAAAA |
| SP2 | GAAGCTTATCGAATTCACTAGTTCAGCCATGATAAACAAGACTG |
| SP3 | AGTCTTGTTTATCATGGCTGAACTAGTGAATTCGATAAGCTTC |
| SP4 | TTTTGTTCTCCTCCTTTTTATGTGATTTCCCCCTTAAAAATA |
| SP5 | GTGAAAAATAAAAGCAACCCCGTGCAAAAAG |
| SP6 | GGGGTTGCTTTTATTTTTCACCGACTCAGTAAGAGC |
| SP7 | GAACAAAATTGTTAAAAACATATGGATCCAAGCTTACTAGTAGT |
| SP8 | GAAGCTTATCGAATTCACTAGTTCACGGCCGTTTGTATAGTTCATC |
| SP9 | GATGAACTATACAAACGGCCGTGAACTAGTGAATTCGATAAGCTTC |
| SP10 | ACTACTAGTAAGCTTGGATCCATATGTTTTTAACAATTTTGTTC |
| SP11 | CTTTTTGCACGGGGTTGCTTTTATTCGGCCGTTTGTATAGTTCATC |
| SP12 | GATGAACTATACAAACGGCCGAATAAAAGCAACCCCGTGCAAAAAG |
| SP13 | AGGCCGCGGATGCATAGGCCTTTAGAGAGAGGAGGTTCTGGC |
| SP14 | TGGGGATCCGCATGCACTAGTTTAGAAGCCTTTTTGACTGTC |
| SP15 | GACAGTCAAAAAGGCTTCTAAACTAGTGCATGCGGATCCCCA |
| SP16 | GCCAGAACCTCCTCTCTCTAAAGGCCTATGCATCCGCGGCCT |
| SP17 | CTACAAAGACGATGACGACAAGTAAACTAGTGCATGCGGATCCC |
| SP18 | GTCGTCATCGTCTTTGTAGTCGAAGCCTTTTTGACTGTCTCT |

| | |
|---|---|
| SP19 | CTTTAAGAAGGAGATATACCATGGCAAAAAGCCAAGCGAAAAAG |
| SP20 | CTCAGTGGTGGTGGTGGTGGTGGGCGGCTTTTTGGGGCACAAAAAAATC |
| SP21 | GATTTTTTTGTGCCCCAAAAAGCCGCCCACCACCACCACCACCACTGAG |
| SP22 | CTTTTTCGCTTGGCTTTTTGCCATGGTATATCTCCTTCTTAAAG |
| SP23 | CCGCGGGGTGCAACAGGATCCGCAAAAAGCCAAGCGAAAAAG |
| SP24 | CTGGTCTGATCGGATCTCTAGGCGGCTTTTTGGGGCACAAAAAAATC |
| SP25 | GATTTTTTTGTGCCCCAAAAAGCCGCCTAGAGATCCGATCAGACCAG |
| SP26 | CTTTTTCGCTTGGCTTTTTGCGGATCCTGTTGCACCCCGCGG |
| SP27 | GATTATACCGAGGTATGAAAACCTTGGTCTGATAATGGGATTTAC |
| SP28 | TCTGTAAAGGTCCAATTCTCGGTGATCACTCCCTTTTTTATTTTC |
| SP29 | CGAGAATTGGACCTTTACAGA |
| SP30 | TTTTCATACCTCGGTATAATC |
| SP31 | TTACTGGATGAATTGTTTTAGCATTCATCTCTATTGTTTCTT |
| SP32 | TTAGACATCTAAATCTAGGTAGTTTTATACATAGAAACAGCA |
| SP33 | TACCTAGATTTAGATGTCTAAAAAGC |
| SP34 | CTAAAACAATTCATCCAGTAA |
| SP35 | TAACTTTAAGAAGGAGATATACCAATGTTAGACCGTTTAAAATCAA |
| SP36 | TTATTAGTGGTGGTGGTGGTGGTGACCTTCCGACTGCTGAAGCTTG |
| SP37 | CACCACCACCACCACCACTAATAATGAGATCCGGCTGCTAACAAAG |
| SP38 | TGGTATATCTCCTTCTTAAAGTTA |
| SP39 | TAACTTTAAGAAGGAGATATACCAATGGAATTATCAGAAATTAGAGC |
| SP40 | TTATTAGTGGTGGTGGTGGTGGTGTGAAAGCTTAGAACGCAGGTAG |
| SP41 | CATACAGCAGTTGATGATAAGCACCACCACCACCACCACTGAG |
| SP42 | CATGAATGGTCTTCGGTTTCCG |
| SP43 | CGGAAACCGAAGACCATTCATG |
| SP44 | CTCAGTGGTGGTGGTGGTGGTGCTTATCATCAACTGCTGTATG |
| SP45 | GGCCTGGTGCCGCGCGGCAGCAGTCGATATCAGCATACTAAA |
| SP46 | GGCTTTGTTAGCAGCCGGATCTTAGTGATTTACTTTCTTGCCACT |
| SP47 | GATCCGGCTGCTAACAAAGCC |
| SP48 | GCTGCCGCGCGGCACCAGGCC |
| SP49 | GTGCGGATCTCACCAACAGATTCATCAGGCCGCCGC |
| SP50 | GCGGCGGCCTGATGAATCTGTTGGTGAGATCCGCACAAG |
| SP51 | GGCCTGGTGCCGCGCGGCAGCGAATTATCAGAAATTAGAGCAG |
| SP52 | AGGTCAAGAGACCCCCTAAAGTCCGC |
| SP53 | AGGGGGTCTCTTGACCTCGAATCAAAGGA |
| SP54 | GGCTTTGTTAGCAGCCGGATCTTATGAAAGCTTAGAACGCAG |

| | |
|---|---|
| SP55 | TATAAAGACGACGACGACAAATAGAGATCCGATCAGACCAGT |
| SP56 | GTCGTCGTCGTCTTTATAGTCGGCGGCTTTTTGGGGCACAAAAAAATC |
| SP57 | GTCGTCGTCGTCTTTATAGTCCTTATCATCAACTGCTGTATG |
| SP58 | ACCATACAGCAGTAGACGACAAAGACTTCTTCGTGCCCCAAAAAG |
| SP59 | TCGTCTACTGCTGTATGGTCGTACGGGTTCTTATGCTTCCGTTTAT |
| SP60 | GCTCGTGTTGTACAATAAATGTAGGAATCCTTAAGGTTTACGGTTTTAGAGCTAGAAATAGC AAGTTAAAATAAGGC |
| SP61 | ACATTTATTGTACAACACGAGCC |
| SP62 | TTCGATAAGCTTCTAGGATCCCATGCAGCTCTTACAGCAGTG |
| SP63 | GGCCAAAAAACTGCTGCCTTCCTAGGCGGCTTTTTGGGGCAC |
| SP64 | GAAGGCAGCAGTTTTTTGGCCTTC |
| SP65 | GGATCCTAGAAGCTTATCGAA |
| SP66 | GCTCGTGTTGTACAATAAATGTGTGTTTACGAGATCGCCTCTGTTTTAGAGCTAGAAATAGCA AGTTAAAATAAGGC |
| SP67 | GGCGCGGGCGCACAGCACGTCAATACGACG |
| SP68 | GCCCGCGCCGCTTGCACGGTA |
| SP69 | GATCCTAGAAGCTTATCGAATTCC |
| SP70 | GCAGTCTAGACTCGAGTAAGG |
| SP71 | CCTTACTCGAGTCTAGACTGCTCGAATTCTCATGTTTGACAG |
| SP72 | GGAATTCGATAAGCTTCTAGGATCCGATCA |
| SP73 | TTAACTAATAAGGAGGACAAACATGTCCAATTTACTGACCGT |
| SP74 | CCGGTTATTATTACTAATCGCCATCTTCCA |
| SP75 | CGATTAGTAATAATAACCGGGCAGGCCATG |
| SP76 | TTGTCCTCCTTATTAGTTAATCAATTCAAGCTTAATTGTTAT |
| SP77 | AAGCTTGGCGTAATCATGGTC |
| SP78 | GAGCTCGAATTCCTGCAGCTG |
| SP79 | GGTACCCGGGGATCCACTAGT |
| SP80 | GCATGCCTCGAGGGGCCGCCC |
| SP81 | CAGCTGCAGGAATTCGAGCTCATCTCACCCGCCACTGCTTTT |
| SP82 | ACTAGTGGATCCCCGGGTACCCACATTTTCACCTTTCTTTGA |
| SP83 | GGGCGGCCCCTCGAGGCATGCTTCTTTGTATCGAATCAGCTT |
| SP84 | GACCATGATTACGCCAAGCTTTTTTCCGGTCCGTTTTGACAG |
| SP85 | GTGAATTCGATAAGCTTCTAGAAGCTAGGAGGAGGATGTGATGACAATGTTTGT |
| SP86 | CACACAAATTAAAAACTGGTCT |
| SP87 | AGACCAGTTTTTAATTTGTGTG |
| SP88 | TAGAAGCTTATCGAATTCAC |
| SP89 | GGTACCCGGGGATCCACTAGT |
| SP90 | GCATGCCTCGAGGGGCCGCCC |

| | |
|---|---|
| SP91 | CAGCTGCAGGAATTCGAGCTCTAAATTGACAATGCAGTCCAG |
| SP92 | ACTAGTGGATCCCCGGGTACCATTCTCCTCCTTGTTCTCTTA |
| SP93 | GGGCGGCCCCTCGAGGCATGCGCTGATGCTCCGCTCGATATG |
| SP94 | GACCATGATTACGCCAAGCTTATTTCCATGCCCATCGCCATC |
| SP95 | TAACTTTAAGAAGGAGGGAGATATACCAATGACAATGTTTGTGGGATC |
| SP96 | TTTGTATAGTTCATCCATGCC |
| SP97 | TTATTATAAAAGAAGAGAACC |
| SP98 | GAAATTAATACGACTCACTATAGGGAGACCACAACGGTTTCCCTCTAGAAATAATTTTGTTTA ACTTTAAGAAGGAG |
| nlpD rev | CGGCGGTCTAATCAACATAC |

### Supplementary Table 6.

| Plasmid name | Combinations of primers and template DNAs for plasmid construction | | | | | |
|---|---|---|---|---|---|---|
| pNAR756 | SP1, 2 | /PY79 DNA | SP3, 4 | /pCH1141 | | |
| pNAR758 | SP5, 6 | /pNAR756 | | | | |
| pNAR778 | SP7, 8 | /pNR1 | SP9, 10 | /pNAR756 | | |
| pNAR780 | SP7, 11 | /pNR1 | SP10, 12 | /pNAR758 | | |
| pNAR809 | SP13, 14 | /PY79 DNA | SP15, 16 | /pLOSS* | | |
| pNAR813 | SP17, 18 | /pNAR809 | | | | |
| pNAR869 | SP19, 20 | /PY79 DNA | SP21, 22 | /pET28b | | |
| pNAR879 | SP23, 24 | /PY79 DNA | SP25, 26 | /pCH913 | | |
| pNAR901 | SP27, 28 | /PY79 DNA | SP29, 30 | /pCH1142 | | |
| pNAR903 | SP31, 32 | /PY79 DNA | SP33, 34 | /pNAR901 | | |
| pNAR913 | SP35, 36 | /PY79 DNA | SP37, 38 | /pET28b | | |
| pNAR915 | SP39, 40 | /pCH2238 | SP37, 38 | /pET28b | | |
| pNAR916 | SP41, 42 | /pNAR869 | SP43, 44 | /pNAR869 | | |
| pNAR917 | SP45, 46 | /JM109 DNA | SP42, 47 | /pET28b | SP43, 48 | /pET28b |
| pNAR939 | SP49, 50 | /pNAR915 | | | | |
| pCH2238 | SP51, 52 | /PY79 DNA | SP53, 54 | /PY79 DNA | SP47, 48 | /pET28b |
| pCH2239 | SP55, 56 | /pNAR879 | | | | |
| pCH2240 | SP55, 57 | /pNAR879 | | | | |
| pCH2241 | SP58, 59 | /pCH2239 | | | | |
| pCH2264 | SP60, 61 | /pKIG855 | | | | |
| pCH2293 | SP62, 63 | /PY79 DNA | SP64, 65 | /pDG1731 | | |
| pCH2302 | SP61, 66 | /pKIG855 | | | | |
| pCH2307 | SP67, 68 | /pNAR915 | | | | |

| | | | | | | | | |
|---|---|---|---|---|---|---|---|---|
| pKIG855 | SP69, 70 | /synthetic DNA | SP71, 72 | /pDG1662 | | | | |
| pMK2 | SP73, 74 | /P1 phage DNA | SP75, 76 | /pLOSS* | | | | |
| pKY13 | SP77, 78 | /pCH1142 | SP79, 80 | /pCH1142 | SP81, 82 | /PY79 DNA | SP86, 87 | /PY79 DNA |
| pNR1 | SP85, 86 | /pCH747 | SP87, 88 | /pCH1130 | | | | |

### Supplementary Table 7. PCR templates and primers for preparation of in vitro translation templates

| Gene name | 1st PCR | | | 2nd PCR | | |
|---|---|---|---|---|---|---|
| | Primers | | template | Primers | | template |
| GFP-ns | SP95 | SP96 | pCH805 | SP98 | SP96 | 1st PCR product |
| GFP-*mifM* | SP95 | SP97 | pCH805 | SP98 | SP97 | 1st PCR product |
